# Increasing heart vascularisation after myocardial infarction using brain natriuretic peptide stimulation of endothelial and WT1^+^ epicardium-derived cells

**DOI:** 10.1101/2020.07.17.207985

**Authors:** Na Li, Stéphanie Rignault-Clerc, Christelle Bielmann, Anne-Charlotte Bon-Mathier, Tamara Déglise, Alexia Carboni, Mégane Ducrest, Nathalie Rosenblatt-Velin

## Abstract

Brain natriuretic peptide (BNP) treatment increases heart function and decreases heart dilation after myocardial infarction. Here, we investigated whether part of the cardioprotective effect of BNP in infarcted hearts related to improved neovascularisation. Infarcted mice were treated with saline or BNP for 10 days. BNP treatment increased vascularisation and the number of endothelial cells in the infarct border and remote zones of infarcted hearts. Endothelial cell lineage tracing showed that BNP directly stimulated the proliferation of resident mature endothelial cells in both areas of the infarcted hearts, via NPR-A binding and p38 MAP kinase activation. BNP also stimulated the proliferation of WT1^+^ epicardium-derived cells but only in the hypoxic area of infarcted hearts. Our results demonstrated that these immature cells have a natural capacity to differentiate into endothelial cells in infarcted hearts. BNP treatment increased their proliferation but not their differentiation capacity. We identified new roles for BNP and new therapeutic strategies to improve heart recovery in infarcted hearts.

## INTRODUCTION

Increased vascularisation supports heart recovery after ischemia. The formation of new vessels in the hypoxic area restores blood flow, provides oxygen and nutriments to the surviving cells, and promotes the migration and engraftment of new cells. While angiogenic inhibition contributes to the development of heart failure in cardiac injury animal models, early heart reperfusion or increased angiogenesis improves cardiac function and delays the onset of heart failure in patients suffering from cardiac ischemia ^1-3^.

Angiogenesis is the main mechanism of neovascularisation in adult infarcted hearts ^4-6^. The origin of new endothelial cells (i.e., resident or infiltrating) as well as the underlying mechanism leading to their proliferation (partial endothelial-to-mesenchymal transition [EndMT] or not) have long been debated. The current consensus is that after myocardial infarction (MI), angiogenesis in the infarct border zone of the heart occurs by clonal expansion of pre-existing resident endothelial cells with no EndMT mechanism ^4, 5, 7^.

Stimulating angiogenesis after MI can improve heart recovery. One complementary therapy could be “re-activating” vasculogenesis (i.e., the differentiation of precursor cells into mature endothelial cells), a mechanism that occurs in the heart during development but is quiescent in adult hearts. Epicardial cells, and more precisely, cells expressing the Wilms’ tumour 1 transcription factor (WT1) migrate from the epicardium to the myocardium during heart development and then differentiate into coronary endothelial cells, pericytes, smooth muscle cells, and even cardiomyocytes after epithelial-to-mesenchymal transition ^8-10^. Consequently, numerous WT1^+^ cells are found in foetal and neonatal hearts, whereas only a few cells express WT1 in adult hearts ^11, 12^.

In hypoxic adult hearts, numerous proliferating WT1^+^ cells can be localised in the epicardium near the infarct and border zone, suggesting that hypoxia stimulates either the proliferation of the remaining WT1^+^ cells or the re-expression of WT1 in cardiac cells ^12-14^. WT1^+^ epicardium-derived cells (EPDCs) remain in a thickened layer on the heart surface without migrating into the myocardium or differentiating into other cell types such as cardiomyocytes or endothelial cells ^15^. Different treatments (e.g., injections of thymosin beta 4 or human amniotic fluid stem cell secretome) fail to induce WT1^+^ cell differentiation into endothelial cells in adult hearts after MI ^13, 14^. Despite enhanced WT1^+^ cell proliferation and higher vessel density in these treated infarcted hearts, no WT1^+^ cell differentiation into endothelial cells has been detected, implying that WT1^+^ EPDCs improve neovascularisation in infarcted hearts via paracrine stimulation.

It is therefore important to identify soluble factors to increase neovascularisation in the heart after MI. For several years, we have studied the role of brain natriuretic peptide (BNP) in the heart during ageing and after ischemic damage ^16^. BNP is a cardiac hormone belonging to the natriuretic peptide family along with atrial natriuretic peptide (ANP) and C-type natriuretic peptide (CNP). It is mainly secreted in the ventricles by cardiomyocytes, fibroblasts, and endothelial and precursor cells ^16, 17^. BNP binds to guanylyl cyclase receptors, NPR-A and NPR-B, thus increasing the intracellular cGMP level ^18^.

BNP is synthesised in the cell cytoplasm as pre-proBNP precursor, cleaved by furin and corin into proBNP peptide, and then into the biologically active carboxy-terminal BNP peptide (active BNP) and the inactive N-terminal fragment (NT-proBNP) ^19, 20^. ProBNP, active BNP, and NT-proBNP peptides are continuously secreted by cardiac cells and ProBNP peptide is the major circulating form of BNP in the plasma of healthy individuals ^21^. O-glycosylated proBNP was also detected in the plasma of patients suffering from heart failure ^22^, and O-glycosylation of proBNP prevents cleavage of this peptide ^22^. Thus, the balance between proBNP and active BNP is impaired in patients with heart diseases, as they have higher levels of inactive proBNP and reduced levels of active BNP. Patients with heart disease therefore have a deficit in functional active BNP ^23^.

BNP secreted by cardiac cells acts on different organs such as the kidneys (modulating sodium and water excretion), vessels (increasing dilation), fat (increasing lipolysis), and pancreas (modulating insulin secretion) ^17, 22^. In the heart, the role of BNP remains unclear, although a majority of cardiac cells (cardiomyocytes, fibroblasts, endothelial cells) express BNP receptors in physiological and pathological states ^16^.

In the last few years, we and others reported that BNP supplementation after ischemic damage promotes the recovery of cardiac function and prevents cardiac remodelling in adult rodent ischemic hearts ^16, 24^. Furthermore, in clinic, a treatment (LCZ696 or Entresto, Novartis) based on inhibition of neprilysin, an enzyme involved in the degradation of the natriuretic peptides, leads to reduced rate of mortality in patients suffering from heart failure with reduced and preserved cardiac contractility or ejection fraction ^25^. Although BNP treatment inhibits fibrosis and protects cardiomyocytes from cell death, the cellular mechanisms responsible for this cardioprotective effect have not been fully elucidated ^16, 17, 24^. Curiously, we observed that BNP stimulates *in vitro* and *in vivo* (i.e., in adult infarcted hearts) the proliferation of cardiac non-myocyte cells (NMCs) expressing the stem cell antigen-1 (Sca-1) ^26^. As Sca-1^+^ cells in adult hearts were reported to be pure endothelial cells ^27^, we questioned whether BNP modulates endothelial cell fate.

Thus, this work aimed to determine whether part of the cardioprotective effect of BNP treatment in infarcted hearts relates to increased neovascularisation, since high BNP levels in plasma are associated with increased collateral development in patients with coronary artery disease ^28^.

## RESULTS

### BNP direct action on cardiac non-myocyte cells after intraperitoneal injections

MI was induced in mice by permanent ligation of the left anterior descending artery. Injection of BNP was immediately performed after surgery and then every 2 days for up to 10 days after surgery (**Fig. 1A**). This protocol from our previous studies demonstrated that BNP injections (first intraventricular and then intraperitoneal) reduced heart remodelling and increased Sca-1^+^ cell numbers ^16, 26^. However, as BNP modulates the function of different organs (kidneys, vessels, pancreas), its effect on hearts could be indirect. Thus, we first aimed to determine whether intraperitoneally injected BNP acted directly or indirectly on cardiac cells.

**Figure 1.**
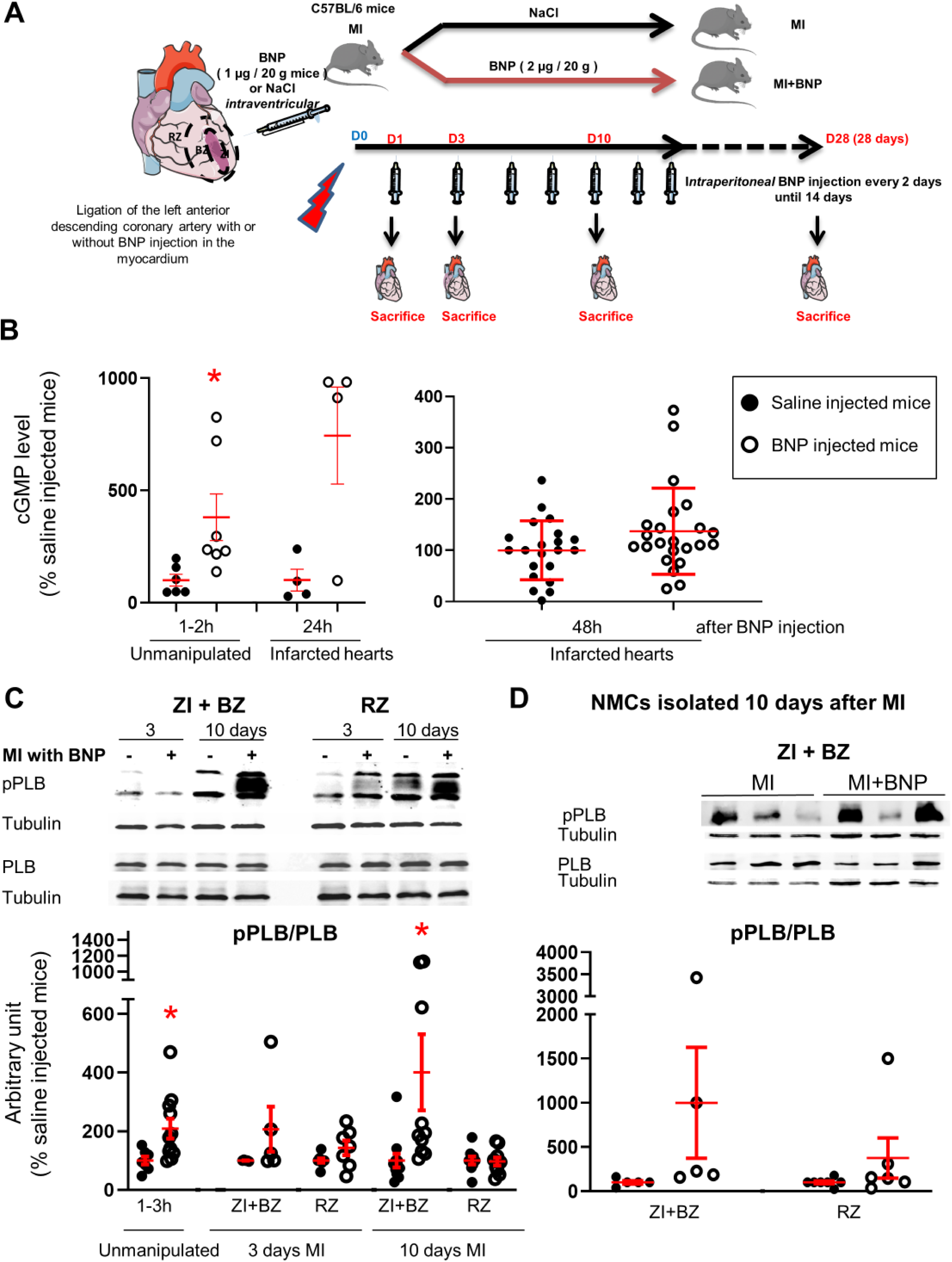
Intraperitoneal BNP injection acts on cardiac non-myocyte cells. **A**. Experimental protocol as described in details in Material and Methods section. B. cGMP plasma level measurement in unmanipulated or infarcted mice injected or not with BNP. n= 6-7 mice for unmanipulated hearts, n= 4 infarcted mice 24h after injection and n= 21-23 infarcted mice 48h after BNP injection. **C**. Representative western blots of total proteins isolated hearts of saline or BNP-injected mice, 3 and 10 days after surgery. Blots were stained with antibodies against phospho phospholamban (pPLB), phospholamban (PBL) and Tubulin (used as loading control). Only the bands at the adequate molecular weight were represented here: Tubulin 55 kDa, pPLB between 17 and 26 kDa and PLB 25 kDa. Quantification of the pPLB/PLB ratio. Data obtained from western blot analysis on unmanipulated (n= 7-11 mice per group) and infarcted hearts of mice treated or not with BNP. Results of BNP treated hearts expressed relatively to the average of saline treated hearts. <u>3 days after MI:</u> n= 5 mice for the ZI+BZ and 7-8 mice for the RZ <u>10 days after MI</u>: n= 10-11 mice for the ZI+BZ and n=9-10 mice for the RZ **D**. Non-myocyte cells (NMCs) were isolated from both areas of infarcted hearts treated or not with BNP 10 days after surgery. Proteins were extracted from these cells (n= 5 independent isolation per group for the ZI+BZ and n=6-7 for the RZ) and pPLB/PLB ratio was evaluated. Only the western blots obtained for NMCs isolated from the ZI+BZ were represented. For **B, C** and **D**: Individual values are represented and the means ± SEM are represented in red. * p<0.05 only for groups with n≥6.

For this purpose, activations of different components of the BNP signalling pathway were evaluated after BNP injections in unmanipulated or infarcted mice (**Fig. 1**). BNP binding to the receptors NPR-A and NPR-B, but not NPR-C, increases intracellular cyclic guanosine 3′,5′-monophosphate (cGMP) levels. cGMP modulates the protein kinase G activity and induces phosphorylation at Ser16 of phospholamban (PLB) ^17^. Increased intracellular cGMP levels simultaneously activate GMP-related cation channels, leading to cGMP extrusion from the cytoplasm into the circulation.

In the plasma of unmanipulated mice, cGMP concentration increased 3.8-fold (103 vs 25 pmoles/ml) after one BNP injection (**Fig. 1B**). In infarcted mice, 1 day after BNP injection, the plasmatic cGMP concentration remained high (265 vs 36 pmoles/ml), but it returned to the level detected in the plasma of saline-injected mice 2 days after BNP injection. The increased plasmatic cGMP concentration demonstrates that injected BNP binds to NPR-A and/or NPR-B receptors.

The increase in PLB phosphorylation after BNP injections was evaluated by western blot analysis on proteins extracted from all cardiac tissue or isolated NMCs (**Fig. 1C-D**). The pPLB/PLB ratio increased 2.1-fold in “unmanipulated” hearts 1 to 3h after BNP injection (**Fig.1C**). In infarcted hearts of BNP-treated mice, the pPLB/PLB ratio increased 3 days after MI, 2.1 and 1.5-fold in the infarct and border zone (ZI+BZ) and in the remote zone (RZ), respectively. 10 days after MI, the pPLB/PLB ratio increased 4.0-fold in the ZI+BZ and was unchanged in the RZ. Furthermore, the pPLB/PLB ratio increased 197- and 179-fold in NMCs isolated from the ZI+BZ and RZ of infarcted hearts in BNP-treated mice 10 days after MI (**Fig. 1D**). According to these results, intraperitoneal injections of BNP can target cardiac NMCs.

### Increased number of endothelial cells in infarcted hearts

NMCs were isolated from both areas of infarcted hearts in saline and BNP-treated mice. We performed quantitative reverse transcription polymerase chain reaction for genes specific to endothelial cells, detecting increased mRNA levels coding for CD31 (x 1.3, p= 0.025) and Ve-cadherin (x 1.3, p= 0.07) in the cells from RZ of BNP-treated infarcted hearts 3 days after MI (**Fig. 2A**). 10 days after MI, mRNA levels coding for vWF (x 1.5, p= 0.023), VeCad (x 1.4, p=0.0007) and eNOS (x 1.4, p=0.049) were increased in the ZI+BZ after BNP treatment (**Fig. 2A**). The number of CD31^+^ cells per mg of cardiac tissue was then evaluated in the ZI+BZ and RZ of infarcted hearts by flow cytometry analysis (**Fig. 2B-C**). The number of CD31^+^ cells increased in the RZ (x 2) 3 days after MI following BNP treatment (**Fig. 2C**). A higher number of CD31^+^ cells was found 10 days after MI in the ZI+BZ (+29%, p= 0.04) and RZ (+28%, p=0.01) of BNP-treated hearts (**Fig. 2C**). This was confirmed by western blot analysis (**Fig. 2D**). CD31 protein levels were higher in the ZI+BZ (+26%, p=0.06) and RZ (+69%, p=0.0003) of BNP-treated hearts compared to saline-injected hearts 10 days after MI (**Fig. 2D**).

**Figure 2.**
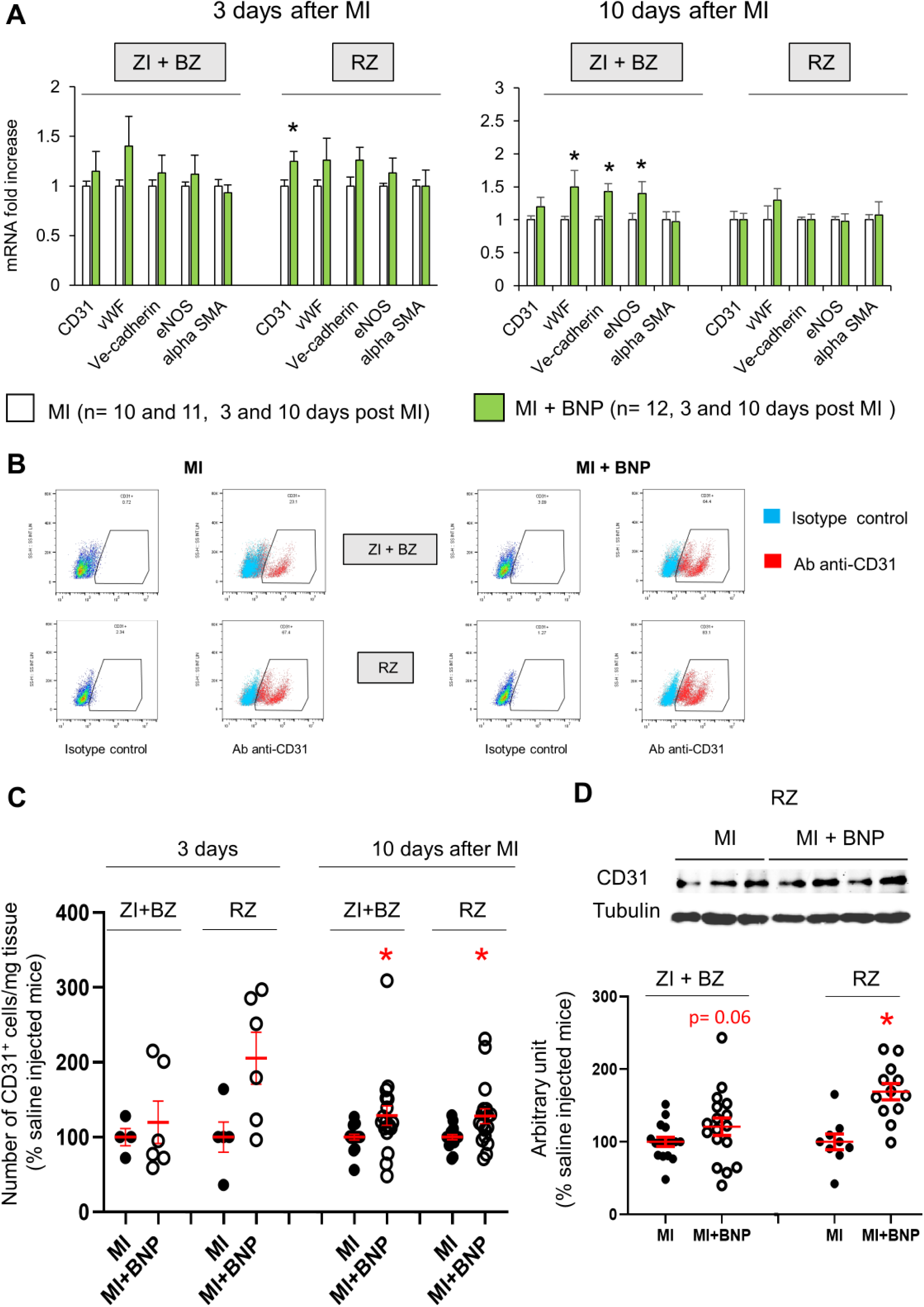
Increased endothelial cell number in infarcted hearts after BNP treatment. **A**. Quantitative relative expression of mRNAs coding for endothelial cell specific genes (CD31, von Willbrand factor (vWF), Ve-cadherin, eNOS, alpha smooth muscle actin (alpha SMA)) in the ZI+BZ and RZ areas of saline (MI) and BNP-injected hearts (MI+BNP) 3 and 10 days after surgery. Results expressed as fold-increase above the levels in saline injected infarcted mice. Results are represented as mean ± SEM. * p<0.05 **B**. Representative flow cytometry analysis of NMCs isolated from the ZI+BZ or RZ of infarcted hearts after BNP or saline treatments 10 days after MI. NMCs stained with control isotype or antibody against CD31 protein. Analysis performed on DAPI negative cells (i.e., living cells). **C**. Quantification of the data obtained by flow cytometry analysis on NMCs isolated from infarcted hearts 3 and 10 days after MI. The number of CD31^+^ cell in BNP treated-hearts related to the number obtained in saline-injected hearts. <u>3 days after MI</u>: n= 4 MI and 6 MI + BNP mice. <u>10 days after MI:</u> n= 16 MI and 15 MI +BNP mice. **D**. Representative western blot of proteins extracted from the ZI+BZ of MI and MI+BNP hearts 10 days after surgery. Blots were stained with antibodies against CD31 and Tubulin (used as loading control). Only the bands at the adequate molecular weight were represented here: Tubulin (55 kDa), CD31 (130 kDa). Quantification of the data from western blot analysis expressed relative to the average of MI hearts. Results were from n= 15-16 different hearts for the ZI+BZ and n=9-12 hearts for the RZ. **C, D:** Individual values are represented and the means ± SEM are represented in red, *p<0.05 only for groups with n≥6.

Finally, cardiac vascularisation (evaluated by CD31 staining intensity) was determined 3, 10, and 28 days after MI in the BNP- or saline-treated hearts of mice (**Fig. 3A-B**). Cardiac vascularisation increased 2.2-fold 3 days after MI in the RZ (p= 0.002) of BNP-treated hearts, while it remained unchanged in the ZI+BZ. BNP treatment increased cardiac vascularisation 10 after MI in the ZI+BZ (+ 108%, p= 0.02) and RZ (+76%, p=0.002) (**Fig. 3A-B**). 4 weeks after MI, vascularization remained 1.7-fold increased in BNP-treated hearts. We counted CD31^+^ cells on heart slices after immunostaining (**Fig. 3C**), observing a 2.0 and 1.8-fold increase 3 days after MI in the ZI+BZ (p= 0.003) and RZ (p= 0.024) of BNP-treated hearts compared to saline-injected hearts, respectively. A 1.4- and 2-fold increase in CD31^+^ cells was counted 10 days after MI in the ZI+BZ (p=0.02) and RZ (p= 0.05) of BNP-treated mice, respectively. This was also the case 28 days after MI (ZI+BZ: x 1.8, and RZ: x 2) (**Fig. 3C**).

**Figure 3.**
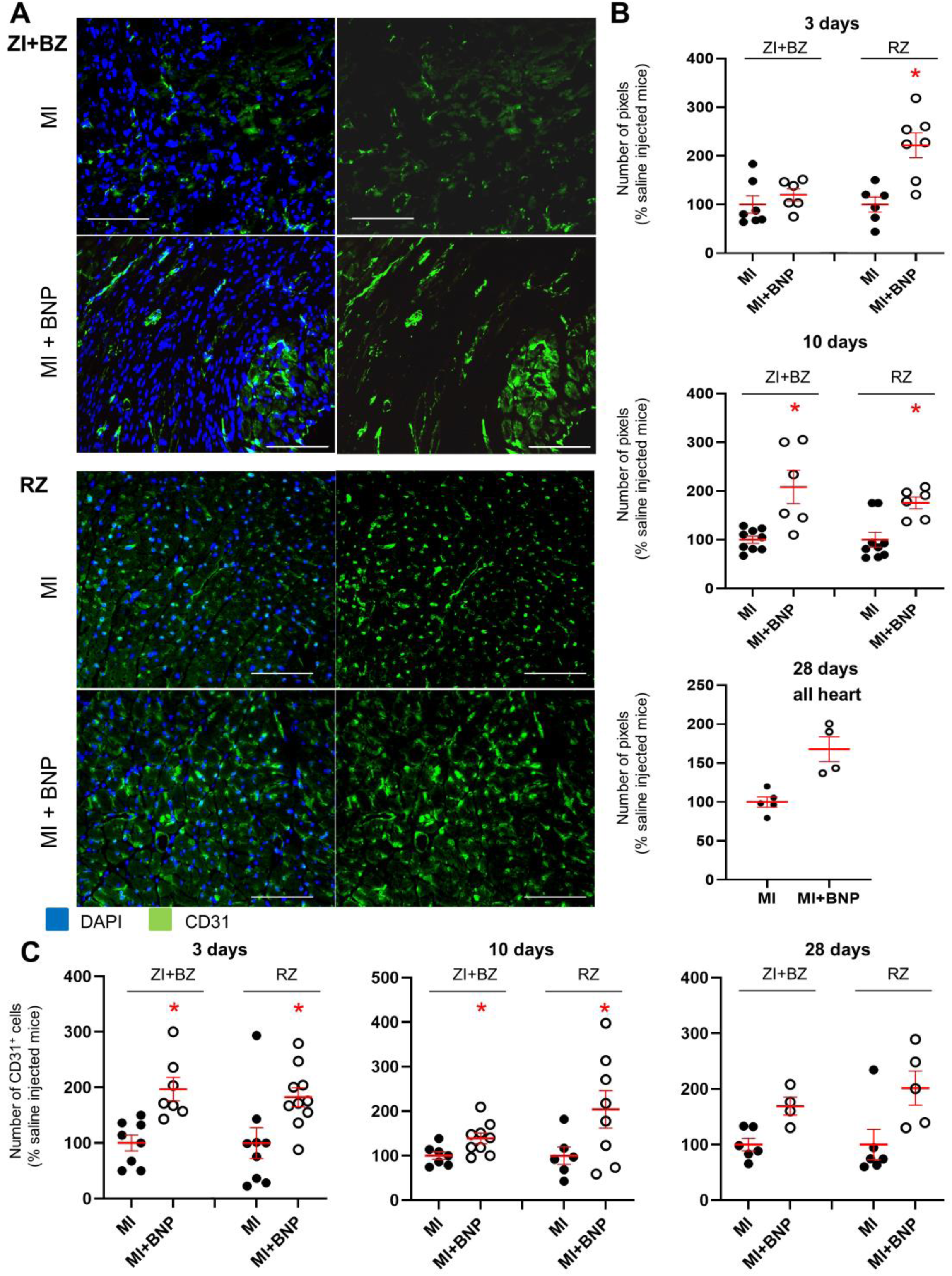
Increased vascularisation in BNP-treated infarcted hearts. **A**. Representative immunostainings against CD31 protein (green) on hearts removed from saline-(MI) and BNP-treated infarcted mice (MI + BNP) 10 days after surgery. Nuclei stained in blue with DAPI. Scale bars: 100 μm **B**. CD31 staining intensity measured on at least 10 different pictures per heart and per area 3, 10 and 28 days after MI. Number of pixel in BNP-injected mice related to the numbers of saline-injected mice. **C**. CD31^+^ cell number counted on heart sections of the different area of saline- and BNP-treated infarcted hearts. Cells counted on at least 10 different pictures per area and mouse. **B-C**: Individual values are represented and the means ± SEM are represented in red. * p<0.05 only for groups with n≥6.

*In vitro* studies allowed identifying by which receptor BNP acts. NMCs isolated from neonatal or adult hearts were treated or not with BNP (**Fig. 4**). mRNA levels coding for von Willbrand factor, Ve-cadherin, and eNOS were upregulated in neonatal BNP-treated cells compared to untreated cells (**Fig. 4A**). BNP treatment increased the number of CD31^+^ cells *in vitro* (+87% and +41% in neonatal and adult NMCs, respectively) (**Fig. 4C**). The increase in CD31^+^ cells after BNP treatment completely disappeared in the NMCs isolated from the NPR-A knockout mouse model (KO) but not from NPR-B KO hearts (+56%, p= 0.04), suggesting that BNP binds to NPR-A to increase the number of endothelial cells (**Fig. 4C**).

**Figure 4.**
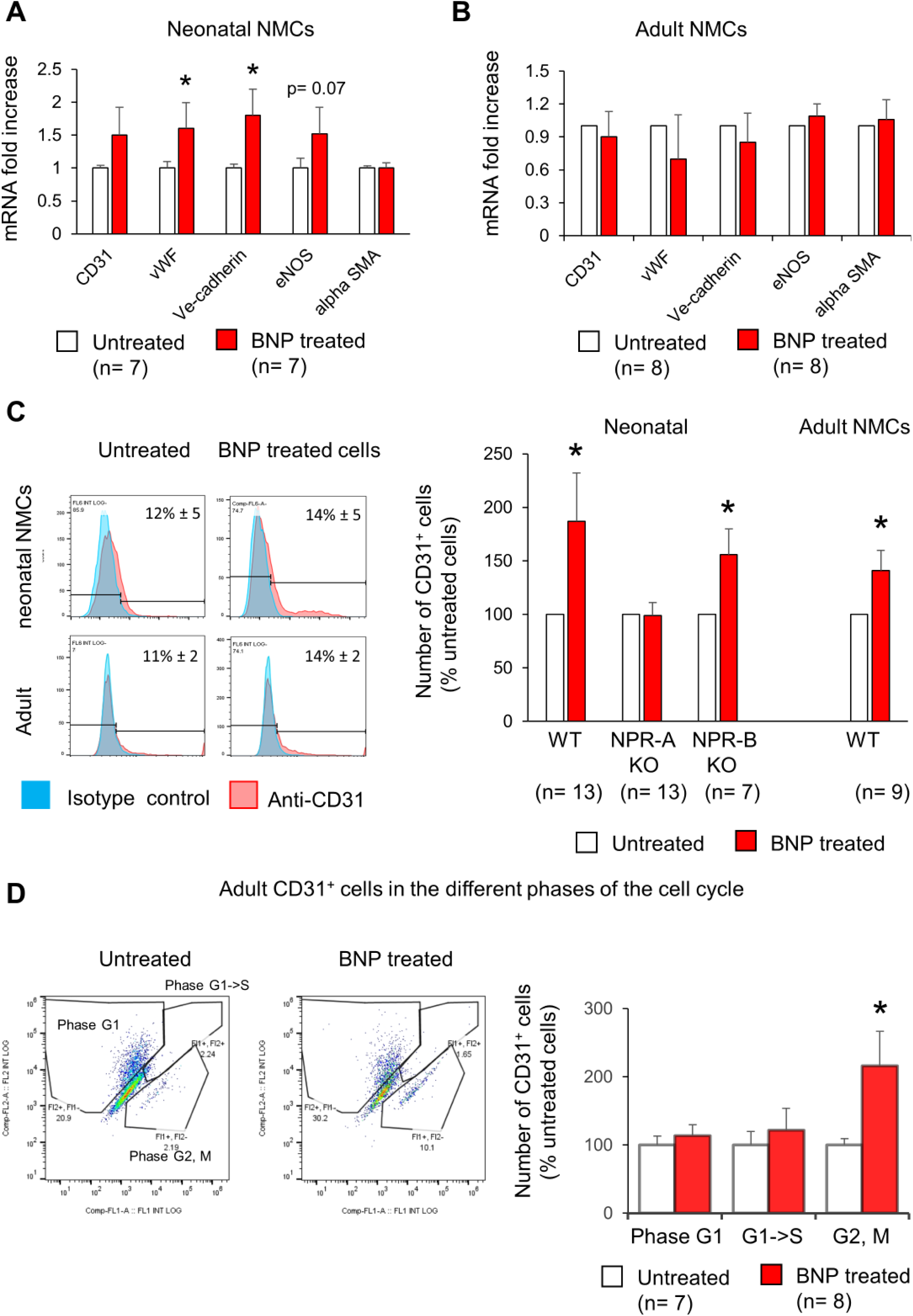
Increased number of endothelial cells *in vitro* after BNP treatment. **A-B**. Quantitative relative expression of mRNAs coding for endothelial cell specific genes (CD31, von Willbrand factor (vWF), Ve-cadherin, eNOS, alpha smooth muscle actin (a SMA)) in NMCs isolated from neonatal (**A**) or adult (**B**) hearts cultured until confluence with or without BNP. Results expressed as fold-increase above the levels in untreated cells. **C**. Flow cytometry analysis to determine the percentage of CD31^+^ cells (**left**: Representative histograms) in untreated or BNP treated NMCs isolated from neonatal or adult hearts. **Right**: Quantification of the number of CD31^+^ cells. Results expressed as fold-increase above the number obtained in untreated cells. Neonatal NMCs were isolated from heart of C57BL/6 (WT), NPR-A or NPR-B deficient pups. Adult NMCs were isolated only from C57BL/6 hearts. **D**. Adult NMCs isolated from FUCCI mice and treated with or without BNP. Flow cytometry analysis (**left**: representative dot plots) to determine among the CD31^+^ cells, the percentages of cells in the G1 phase (Fl2+ Fl1-), in the G1->S phase (Fl2+Fl1+) and in the G2, M phase (Fl2-Fl1+). **Right:** Quantification of the number of CD31^+^ cells in the different phases of the cell cycle. Results expressed as fold-increase above the numbers obtained in untreated cells. For all quantifications, the results are means ± SEM, * p<0.05

Overall, our results clearly demonstrate that BNP injections after MI lead to more endothelial cells initially in the RZ (3 days after MI) and then in the ZI+BZ (10 days after MI). This is also the case in unmanipulated adult hearts where intraperitoneal BNP injections every 2 days for 10 days increase heart vascularisation (+37%, p=0.0007) (**Supplementary Fig. 1**). *In vitro* experiments suggest that BNP acts via NPR-A.

### Mobilisation of resident mature endothelial cells and precursor cells

The increased number of endothelial cells in infarcted hearts after BNP treatment resulted from either the direct effect of BNP on pre-existing cardiac endothelial cells (i.e., endothelial cells in infarcted hearts express NPR-A and NPR-B (**Supplementary Fig. 2**)), and/or the effect of BNP on other cells. Indeed, BNP may increase the number of infiltrating endothelial cells or stimulate the differentiation of endothelial precursor cells into endothelial cells in infarcted hearts. We therefore studied the origin of endothelial cells in infarcted hearts of BNP-treated mice.

We first investigated whether the increased number of endothelial cells in BNP-treated hearts originated from infiltrating cells. We counted CD31^+^ and CD45^+^ cells in infarcted hearts treated or not with BNP 3 and 10 days after MI (**Supplementary Fig. 3**). The percentage of CD45^+^ cells among the CD31^+^ cell subset was less than 10% in all zones of the infarcted hearts 3 and 10 days after MI. The numbers and percentages of CD45^+^ and CD31^+^ cells were similar in BNP- and saline-treated hearts 3 and 10 days after MI. The increased number of CD31^+^ cells in BNP-treated hearts was thus not due to infiltrating cells.

To understand whether the increased number of endothelial cells in BNP-treated hearts originated from pre-existing endothelial cells or from the differentiation of cardiac precursor cells, heterozygous tamoxifen-inducible Cdh5:ROSA26 mice were used to trace CD31^+^ cells (**Supplementary Fig. 4**). Tamoxifen injections given 2 weeks before MI induced green fluorescent protein (GFP) expression in CD31^+^ cells (**Supplementary Fig. 4 A-C**). To avoid contamination with GFP^-^ cells, our analysis focussed on the CD31^+^ cell subset with 94% expressing the GFP protein before MI (**Supplementary Fig. 4C**). Ten days after MI, almost all CD31^+^ cells expressed the GFP protein in the ZI+BZ and RZ of BNP-treated and untreated infarcted hearts (**Fig. 5A-B**). As shown in **Supplementary Figure 4D**, more GFP^+^ cells were apparent in the ZI+BZ of BNP-treated mice compared to those injected with saline. Numerous vessels and capillaries formed by GFP^+^ cells were detected in this area after BNP injections. However, we also detected some GFP^-^ endothelial cells in the ZI+BZ of BNP- and saline-treated hearts (**Fig. 5B**).

**Figure 5.**
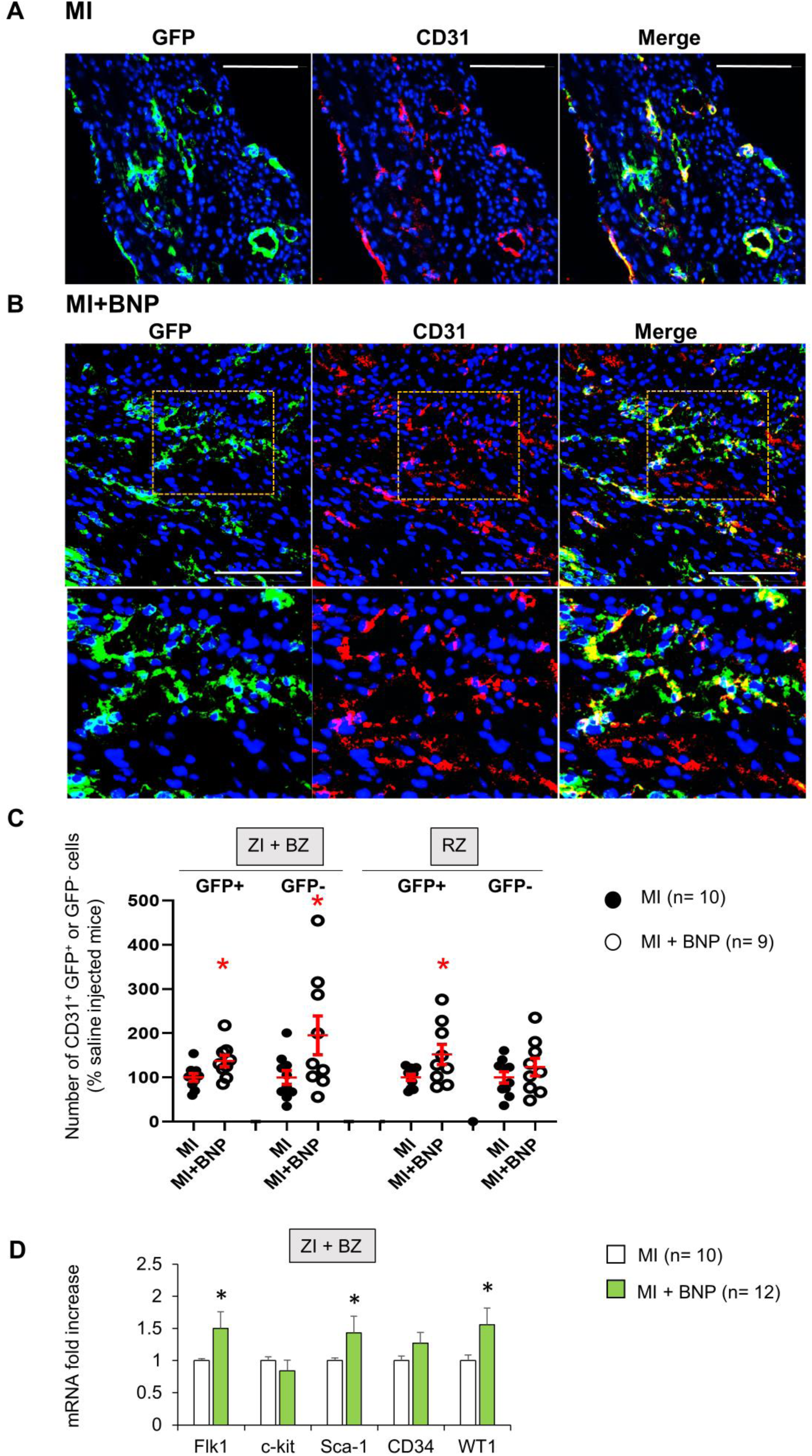
Mobilisation of resident mature endothelial and precursor cells in BNP-treated hearts. **A**. Representative pictures of the ZI+BZ area of Cdh5:ROSA26 infarcted hearts 10 days after surgery, treated (**B**) or not (**A**) with BNP and stained with DAPI (nuclei in blue) and antibody against CD31 protein (red). Endothelial cells or cells originating from CD31^+^ cells express GFP protein. Scale bars represent 100 μm. **B**. Orange rectangles are represented at high magnification below. **C**. Quantification of the number of GFP^+^ and GFP^-^ CD31^+^ cells. Results expressed in BNP-injected mice as fold-increase above the numbers obtained in saline-injected mice. Individual values are represented and the means ± SEM are represented in red. **D**. Quantitative relative expression of mRNAs coding for endothelial precursor specific genes (Flk-1, c-kit, stem cell antigen 1 (Sca-1), CD34 and Wilms’ tumor 1 (WT1)) in ZI+BZ 3 days after MI. Results expressed as fold-increase above the levels in saline-injected mice. Results are means ± SEM **<u>C-D</u>**: * p≤0.05.

We quantified this observation by isolating NMCs in infarcted Cdh5:ROSA hearts. The percentage of CD31^+^ cells expressing the GFP protein was determined by flow cytometry. BNP treatment increased the number of CD31^+^ GFP^+^ cells (i.e., originating from pre-existing endothelial cells) in the ZI+BZ (+37%, p= 0.02) and RZ (+52%, p=0.03) (**Fig. 5C**) of all treated hearts. Interestingly, the number of CD31^+^ GFP^-^ cells increased significantly in the ZI+BZ (+95%, p= 0.05) but not in the RZ (+23%, p=0.2) after BNP treatment (**Fig. 5C**).

Our results demonstrated that endothelial cells in the infarcted hearts of BNP-treated mice originate mainly from pre-existing endothelial cells in the ZI+BZ and RZ. However, endothelial cells originating from precursor cells (i.e., GFP^-^) also contribute to the neovascularisation of the ZI+BZ of BNP-treated infarcted hearts.

### Stimulated proliferation of endothelial cells via p38 MAP kinase activation

We then investigated the capacity of BNP to stimulate the proliferation of endothelial cells (**Fig. 6**) by performing immunostaining against CD31 and 5-Bromo-2′-deoxyuridine (BrdU) on BNP- and saline-treated infarcted hearts 1, 3, and 10 days after surgery (**Fig. 6A-C**). To obtain the percentage of proliferating endothelial cells in each area of the infarcted hearts, the number of CD31^+^ BrdU^+^ cells was divided by the total number of CD31^+^ cells (**Fig. 6C**). During the first 3 days after surgery, 21% of CD31^+^ cells proliferated in the RZ of BNP-treated infarcted hearts versus 14% in the RZ of saline-treated hearts (+53%, p=0.02). In the ZI+BZ, higher endothelial proliferation following BNP treatment was detected only 10 days after MI (+56%, p=0.02) (**Fig.6C**).

**Figure 6.**
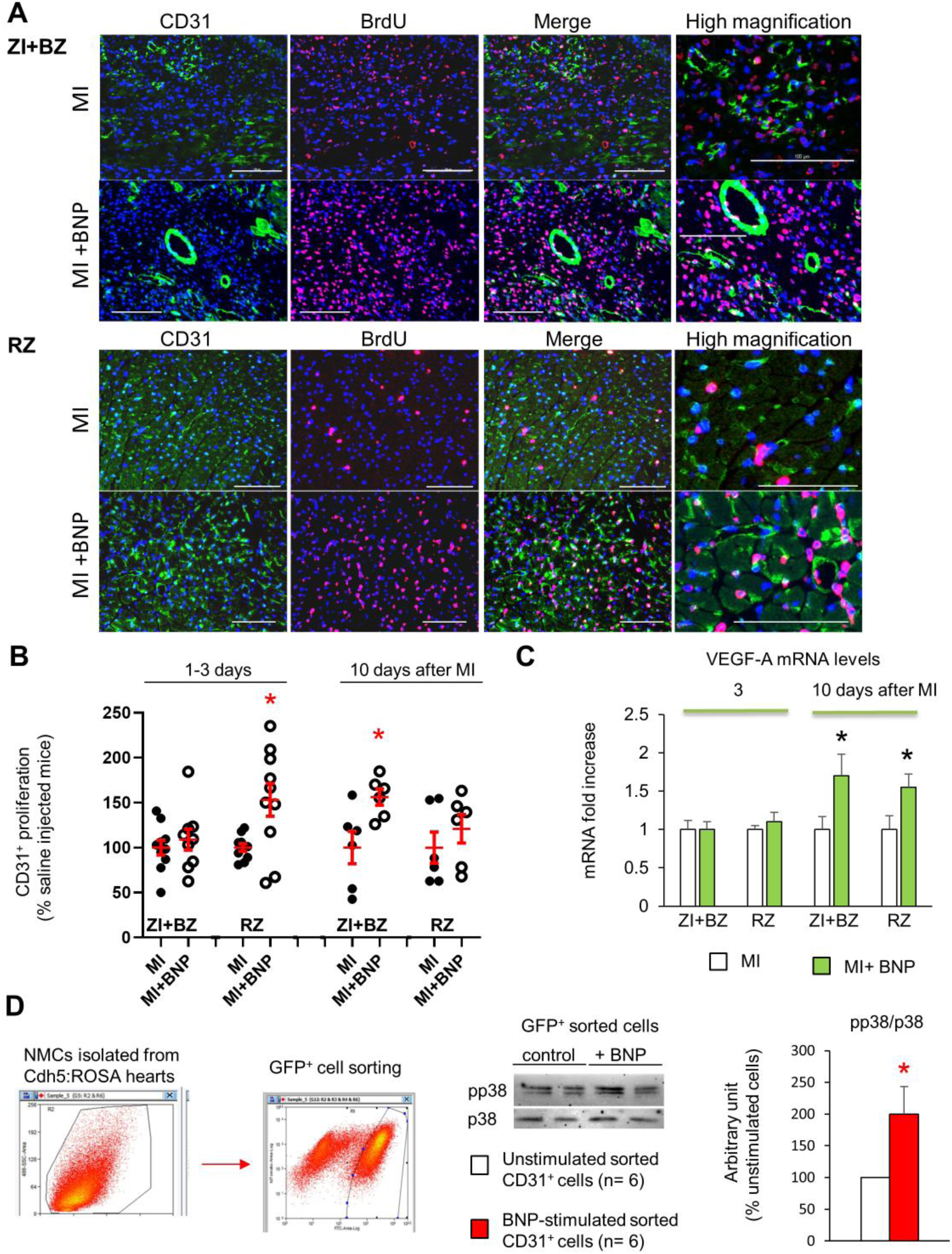
BNP-stimulation of endothelial cell proliferation. **A**. Representative pictures of the ZI+BZ and RZ of C57BL/6 infarcted hearts, 10 days after surgery, treated or not with BNP and stained with DAPI (nuclei in blue) and antibodies against CD31 protein (green) and BrdU (pink). Scale bars: 100 μm. **B**. Percentage of proliferating endothelial cells/per pictures in each area of the infarcted hearts (nber of CD31^+^BrdU^+^ cells / CD31^+^ cells). Results in BNP treated hearts related to those obtained in saline-treated hearts. At least 10 different pictures evaluated per mouse and per area. n= 10-9 mice per group 1-3 days after MI, n=6 mice per group 10 days after MI. Individual values are represented and the means ± SEM are represented in red. **C**. Quantitative relative expression of mRNA coding for VEGF-A in ZI+BZ and RZ 3 and 10 days after MI. Results expressed as fold-increase above the levels in the hearts of saline-injected mice. n= 11-14 hearts per group. **D**. NMCs isolated from unmanipulated Cdh5:ROSA26 mice injected 2 weeks before with tamoxifen. GFP^+^ cells were sorted and stimulated immediately with or without BNP during 1h30 at room temperature. Western bot analysis were then performed on these cells to evaluate p38 MAP kinase activation. Blots were stained with antibodies against phospho p38 (pp38) (43 kDa), p38 (43 kDa) and Tubulin (55 kDa). Quantification of the pp38/p38 ratio obtained from 6 independent cell sorting experiments. **C, D:** Results are means ± SEM, * p<0.05.

To evaluate the direct BNP effect on endothelial cell proliferation, NMCs from adult hearts expressing fluorescent ubiquitination-based cell cycle indicator (FUCCI) were isolated and cultured for 3-4 days with and without BNP (**Fig. 4D**). Transgenic FUCCI mice allow the differentiation of cells in different phases of the cell cycle ^29^. We evaluated the number of CD31^+^ cells in the phases of the cell cycle by flow cytometry after CD31 staining. The number of adult endothelial cells in the G2/M phase of the cell cycle was 2.2 times higher (p=0.04) in BNP-treated compared to untreated NMCs.

To determine whether higher levels of angiogenic factors could be responsible for increased endothelial cell proliferation in BNP-treated hearts, we determined the vascular endothelial growth factor (VEGF)-A mRNA levels in NMCs 3 and 10 days after MI. VEGF-A mRNA levels were similar in NMCs isolated from saline- and BNP-treated hearts 3 days after MI **(Fig.6D**). However, VEGF-A mRNA levels increased 10 days after MI in NMCs isolated from the ZI+BZ (+69%, p-=0.02) and RZ (+55%, p=0.03) of BNP-treated hearts compared to NMCs isolated from saline-treated hearts (**Fig. 6D**).

We determined the signalling pathway activated by BNP on endothelial cells. For this purpose, we sorted GFP^+^ endothelial cells from the hearts of unmanipulated Cdh5:ROSA mice injected with tamoxifen 2 weeks prior (**Fig. 6E**) and then stimulated them for 1.5h with BNP *in vitro*. We extracted proteins from these cells and performed western blot analysis. The pp38/p38 ratio was 2 fold higher after BNP stimulation on the sorted pure endothelial cells compared to untreated cells (p= 0.026). This experiment demonstrated that BNP acts directly on endothelial cells via p38 MAP kinase activation.

### More non-myocyte cells expressing Wilms’ tumour 1 protein

The number of endothelial cells originating from GFP^-^ cells increased in the hypoxic area of hearts isolated from BNP-treated mice, which could point to the mobilisation of vascular precursors by BNP treatment. We measured mRNA levels coding for genes expressed by endothelial precursors in NMCs from the ZI+BZ of BNP-treated or untreated infarcted hearts (**Fig. 5D**). Three days after MI, mRNA levels coding for Flk1 (x 1.5, p=0.01), Sca-1 (x 1.4, p=0.04) and WT1 (x 1.6, p=0.01) increased in cells from the ZI+BZ of BNP-treated infarcted hearts. mRNA levels coding for c-kit did not differ. *In vitro*, NMCs isolated from neonatal hearts were stimulated or not with BNP for 7-10 days. mRNA levels coding for c-kit (x 2.1), Flk1 (x 1.9), Sca-1 (x 2), and WT1 (x 1.6) were significantly higher after BNP treatment (**Supplementary Fig. 5C**). These results suggest that BNP could act on WT1^+^ cells *in vivo* and *in vitro*. We thus verified whether WT1^+^ cells express BNP receptors, NPR-A and/or NPR-B (**Supplementary Fig. 5A**).

We performed immunostaining on BNP- and saline-treated infarcted hearts to evaluate the number of WT1^+^ cells (**Fig. 7A-C**). WT1^+^ cells were easily detected in the epicardium and endocardium of adult hearts after MI as reported by others ^12-14^. In the ZI+BZ area, compared to saline-injected infarcted hearts, BNP treatment led to more WT1^+^ cells 3 days (x 2.5 in epicardium and x 3.5 in endocardium) and 10 days after MI (x 2.9 in epicardium and x 1.7 in endocardium) (**Fig. 7A-C**). In the RZ of BNP-versus saline-treated hearts, the number of WT1^+^ cells increased in the epicardium (x 2.5) and in the endocardium (x 2.3) 3 days after MI and in the epicardium 10 days after MI (x 3.6). No difference in the number of WT1^+^ cells was detected 28 days after MI (data not shown).

**Figure 7.**
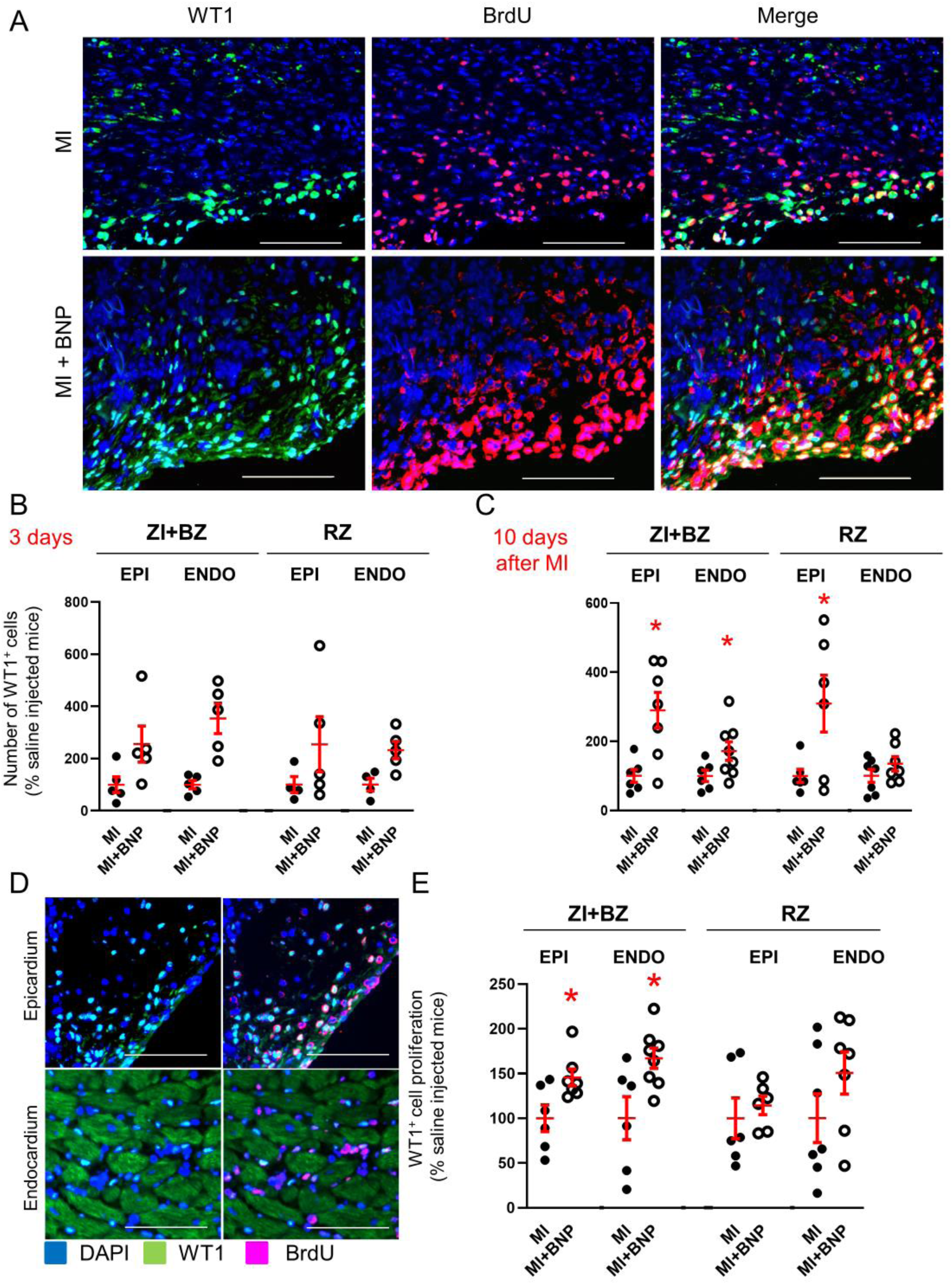
Increased number of WT1^+^ cells in infarcted BNP-treated hearts. **A**. Representative pictures of the epicardium of the ZI+BZ area of C57BL/6 infarcted hearts treated or not with BNP 10 days after MI and stained with DAPI (nuclei in blue) and antibodies against WT1 protein (green) and BrdU (red). Scale bars: 100 μm. **B-C**. WT1^+^ cell number per pictures in the ZI+BZ and RZ of infarcted hearts treated or not with BNP 3 (**B**) and 10 (**C**) days after surgery. **D:** Representative immunostainings of proliferating WT1^+^ cells in the epicardium and endocardium of the ZI+BZ area of BNP treated infarcted heart 10 days after MI. Scale bars: 100 μm. **E**. Percentages of proliferating WT1^+^ cells (nber of WT1^+^BrdU^+^ cells / nber of WT1^+^ cells) 10 days after MI. **B, C, E:** Results obtained in the epicardium separated from those obtained in the endocardium. Individual values are represented and the means ± SEM are represented in red. * p≤0.05 only for groups with n≥6.

### Stimulation of WT1^+^ cell proliferation near the infarct zone

WT1 is re-expressed by mature endothelial cells after hypoxia ^12^. To determine whether BNP stimulates WT1^+^ cell proliferation and/or WT1 re-expression in endothelial cells, the percentage of proliferating WT1^+^ cells (number of WT1^+^ BrdU^+^ cells relative to the total number of WT1^+^ cells) was assessed 3 and 10 days after MI in hearts from BNP-treated or untreated mice.

No increased WT1^+^ cell proliferation was detected in BNP treated hearts 3 days after MI (data not shown). The proliferation of WT1^+^ cells only increased in the epicardium and endocardium of the ZI+BZ in BNP-treated infarcted hearts 10 days after surgery (+45%, p=0.03 and +67%, p=0.04, respectively) (**Fig. 7E**), showing that the higher number of WT1^+^ cells in other areas of BNP-treated hearts was likely due to WT1 re-expression. Interestingly, in BNP treated hearts, almost all WT1^+^ cells localised in the epicardium express BrdU, whereas only 50% of the WT1^+^ cells proliferate in the endocardium (**Fig. 7D**).

Stimulation of WT1^+^ cell proliferation by BNP treatment was also highlighted *in vitro* (**Supplementary Fig. 5D-E**). After BNP stimulation, the number of WT1^+^ cells increased (+38%, p= 0.02) in cultured NMCs isolated from neonatal hearts. BNP treatment stimulated their proliferation (+23%, p= 0.003) compared to untreated NMCs.

### Stimulation of WT1^+^ precursor cell proliferation

The next step was to identify proliferating WT1^+^ cell origin. In order to discriminate between WT1^+^ endothelial precursor cells and mature cells re-expressing WT1 after hypoxia, MI was induced in heterozygous inducible WT1:ROSA mice. Three injections of tamoxifen were administered 2 weeks before MI induction. GFP was expressed only in WT1^+^ cells. Thus, before surgery, 0.4 ± 0.08% of the NMCs were GFP^+^, while 0.6 ± 0.09% of the CD31^+^ cells expressed the GFP protein (n=4 mice).

Ten days after MI, BNP treatment significantly increased the number of GFP^+^ cells by 2.3-fold in the ZI+BZ (p=0.05) of infarcted WT1:ROSA hearts (**Fig. 8A and D**). GFP^+^ cells were mainly localised in the epicardium of the ZI+BZ of infarcted hearts (**Fig. 8A-B (left**). However, in the ZI+BZ of BNP-treated hearts, GFP^+^ cells migrated into the tissue (**Fig. 8A-B (right)**), forming vessel-like structures (**white arrows in Fig. 8A and 8C**). This is confirmed by immunostaining against CD31 (**Fig. 8B-C**). Indeed, some of the GFP^+^ cells stained positive for BrdU, Sca-1, and CD31, showing that WT1^+^ precursor cells can proliferate and differentiate into endothelial cells, especially after BNP treatment (**Supplementary Fig. 6, Fig. 8B (right) and 8C**). In the ZI+BZ of BNP-treated hearts, 10 days after MI, 10.5 ± 2% of CD31^+^ cells expressed the GFP protein versus 5.0 ± 1.3% of endothelial cells in saline-treated hearts (p= 0.03). In the RZ, 8.5 ± 2% of endothelial cells originated from GFP^+^ cells following BNP treatment versus 4.0 ± 1% in saline-treated hearts (p= 0.03) (**Fig. 8E**). However, among the GFP^+^ cells, the percentage of cells differentiating into CD31^+^ cells was the same (around 48%) in the ZI+BZ and RZ between BNP- and saline-treated hearts (**Fig. 8F**).

**Figure 8.**
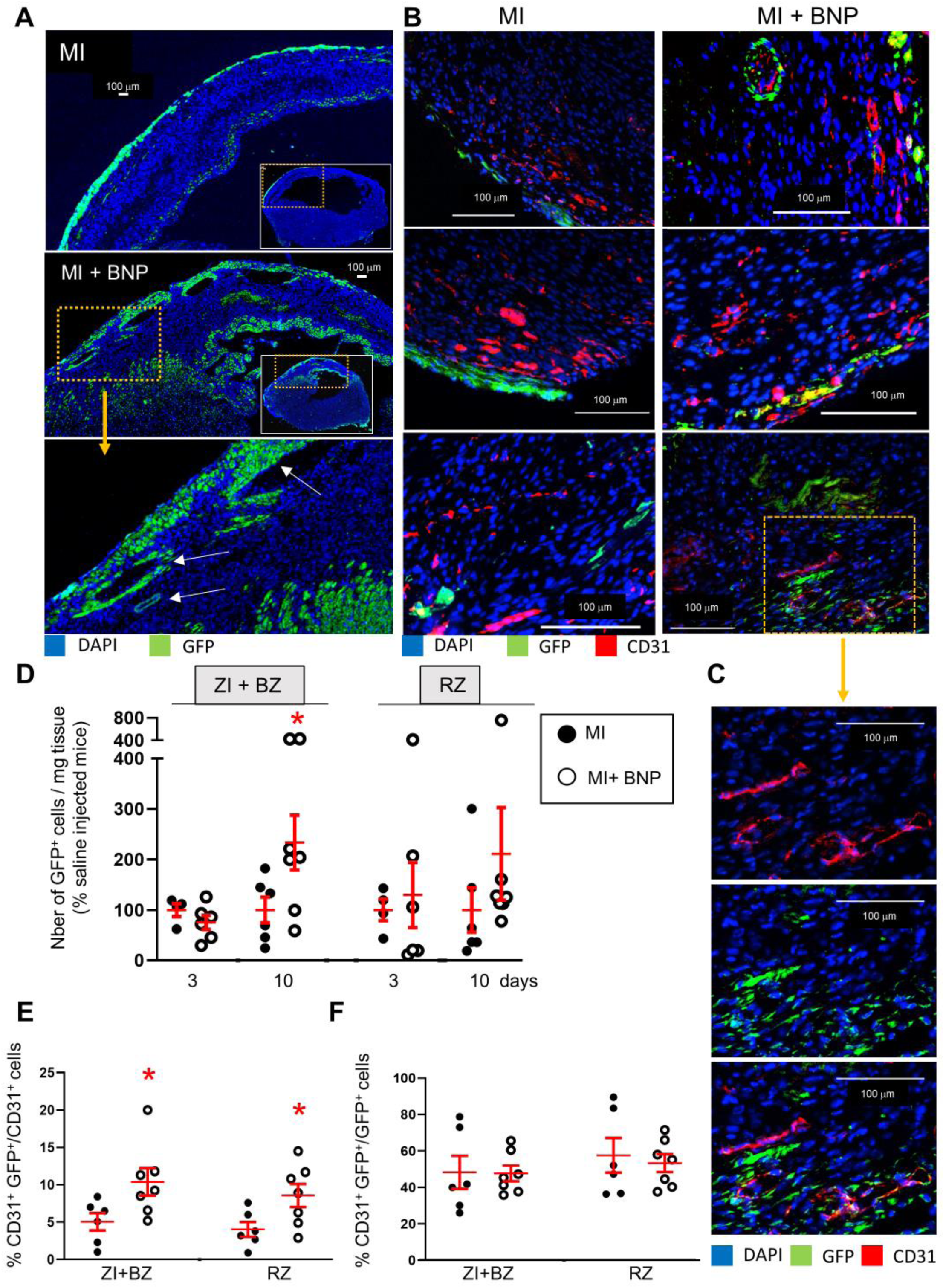
Increased WT1^+^ cell proliferation after BNP treatment in infarcted hearts. **A**. Representative immunostainings of ZI+BZ of WT1:ROSA hearts treated or not with BNP 10 days after surgery and stained with DAPI (nuclei in blue) and antibody against GFP protein (green). Hearts represented in full in the small inserts. The orange rectangles delimited the enlarged area below. **B**. Representative immunostainings of WT1:ROSA hearts treated or not with BNP 10 days after MI and stained with DAPI (nuclei in blue) and antibodies against CD31 protein (red) and GFP (green). White arrows represented GFP^+^ CD31^+^ cells, i.e., endothelial cells originating from WT1^+^ cells. **C**. High magnification of a part of the ZI+BZ of infarcted BNP-treated hearts where WT1^+^ cells contributed to the vessel formation (orange rectangle). **D**. GFP^+^ cell number per mg of cardiac tissue 3 or 10 days after surgery, determined by flow cytometry analysis. Results in BNP treated hearts related to those obtained in saline-treated hearts **E and F**. Flow cytometry analysis on isolated NMCs stained with antibodies against CD31 and GFP. **E**. Percentages of CD31^+^ cells originating from WT1^+^ precursor cells (GFP^+^CD31^+^ cells). The percentages of GFP^+^ cells determined among the selected CD31^+^ cells. **F**. Percentages of differentiating WT1^+^ cells into CD31^+^ cells. The percentages of CD31^+^ cells determined among the selected GFP^+^ cells. **D:** <u>3 days after surgery</u>: MI: n= 4, MI+BNP: n=6 **D-F:** <u>10 days after surgery:</u> MI: n= 6, MI+BNP: n=7 different mice. Individual values are represented and the means ± SEM are represented in red. * p≤0.05 only for groups with n≥6.

Our results demonstrated that WT1^+^ precursor cells have the capacity to differentiate into endothelial cells in infarcted hearts. BNP increased the number of endothelial cells originating from WT1^+^ cells by stimulating WT1^+^ cell proliferation but not their differentiation into endothelial cells.

### Increased vascularization in infarcted hearts after LCZ696 treatment

LCZ696 (Entresto, Novartis) product associates both an angiotensin receptor blocker (valsartan) and an inhibitor of neprilysin (NEP, sacubitril). In the large, randomized, double-blind PARADIGM-HF trial, LCZ696 treatment has been shown to promote significant benefits in patients with chronic heart failure, when compared to angiotensin-converting enzyme inhibition (enalapril) ^25^. NEP is an endopeptidase able to degrade several factors including the natriuretic peptides. Thus, treatments of rats, rabbits and humans with NEP inhibitor increases the blood level of the natriuretic peptides (ANP and BNP) and of cGMP ^30-32^. In the plasma of unmanipulated mice, we determined that cGMP concentration increased 3-fold (138 vs 44.5 pmoles/ml) 24h after LCZ696 treatment (60 mg/kg/day). We thus evaluated LCZ696 treatment on heart neovascularization 10 days after MI. Mice were treated orally by two different concentrations of LCZ696 24h after myocardial infarction induction (**Fig. 9A**).

**Figure 9.**
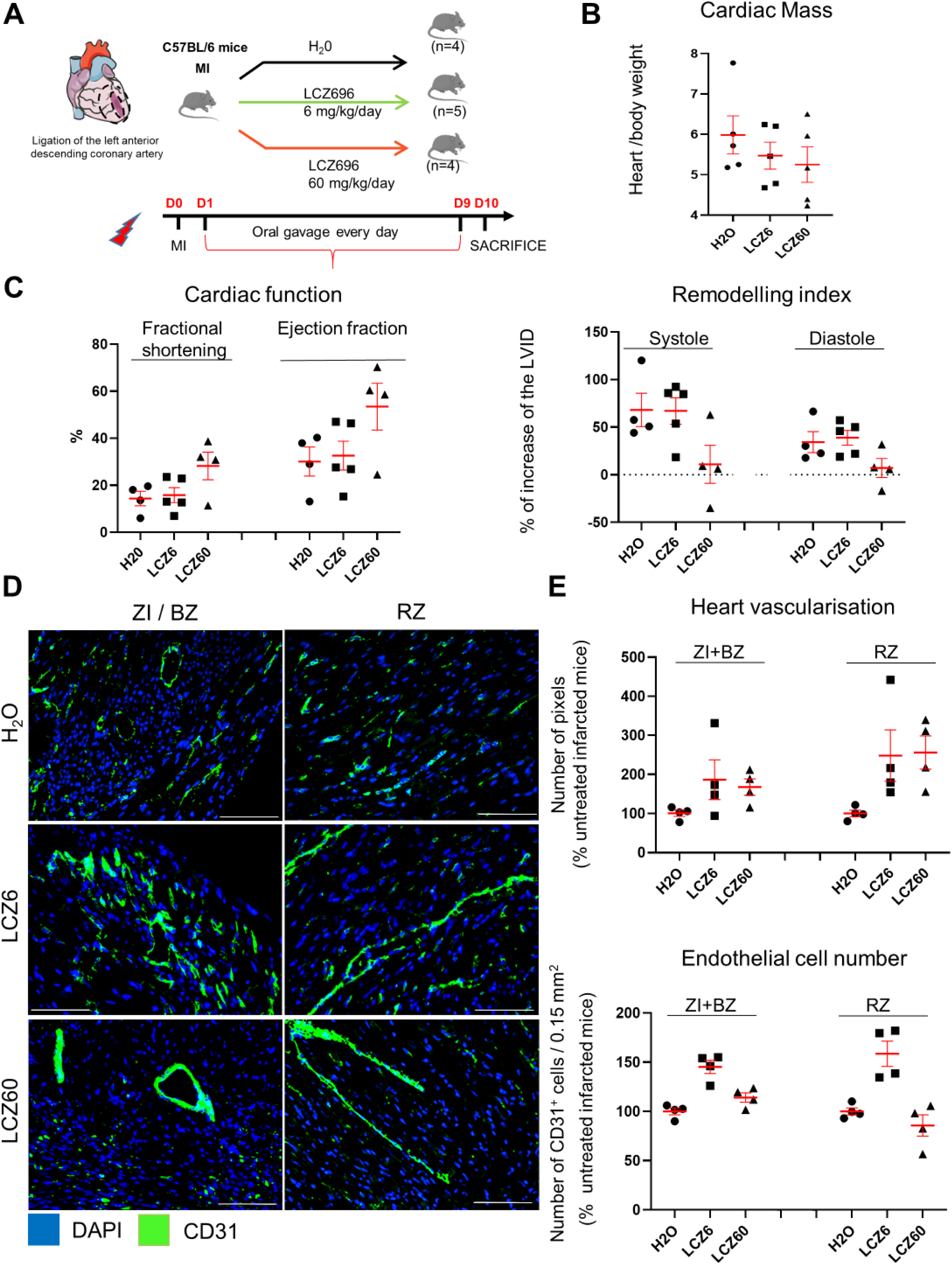
Increased vascularisation in infarcted hearts after LCZ696 treatment. **A**. Experimental protocol as described in details in Material and Methods section. **B**. Cardiac mass (heart weight (mg)/body weight (g)) of infarcted mice 10 days after MI. **C**. Cardiac function and remodelling index measured by echocardiography 10 days after MI before sacrifice. **D**. Representative pictures of the ZI+BZ and RZ areas of infarcted hearts 10 days after surgery, treated with LCZ696 (6 or 60 mg/kg/day) or not and stained with DAPI (nuclei in blue) and antibody against CD31 protein (green). Scale bars represent 100 μm. **E**. CD31 staining intensity measured on at least 10 different pictures per heart and per area 10 days after MI. Number of pixels in hearts of LCZ696 treated mice related to the numbers of untreated mice (H_2_0). CD31^+^ cell number counted on heart sections. Cells counted on at least 10 different pictures per area and mouse. **B, C, E:** Individual values are represented and the means ± SEM are represented in red. No statistical analysis are performed as the number of mice in each group is < 6.

LCZ696 treatment (at the both concentrations) did not affect body weight nor blood pressure. Urea and creatinin plasma levels, were not changed after LCZ696 treatment, demonstrating no altered kidney functions (**data not shown**). LCZ696 treatment prevented the increase of cardiac mass induced by myocardial infarction (−12% at the high concentration) (**Fig. 9B**). Mice treated with high dose of LCZ696 (60 mg/kg/day) displayed increased fractional shortening (+14%) and ejection fraction (+23%) compared to infarcted untreated mice. Moreover, heart remodeling was 6-fold decreased during systole (**Fig. 9C**).

Cardiac vascularisation (evaluated by CD31 staining intensity) increased 1.6-fold in the ZI+BZ and 2.6-fold in the RZ of LCZ696-treated hearts at the both concentrations (**Fig. 9D-E**). A 2.0 - and 2.4-fold increase in CD31^+^ cells was counted in the ZI+BZ and RZ, respectively, of LCZ696 (6 mg/kg/day)-treated hearts (**Fig. 9D-E**). The number of CD31^+^ cells was only slightly increased in the ZI+BZ (x 1.2) and RZ (x 1.3) of high dose of LCZ696 (60mg/kg/day)-treated hearts. Our results demonstrated that LCZ696 administration at both concentrations increased cardiac vascularization in both areas of infarcted hearts. In mice, only the high concentration of LCZ696 was associated with increased heart function and decreased heart remodelling 10 days after MI.

## DISCUSSION

The work presented here continues our previous research aimed at determining the role of BNP in the heart. We already demonstrated that BNP injections in mice after MI decreased heart remodelling and increased heart function ^16^. We thus questioned whether increased neovascularisation in infarcted hearts is part of the cardioprotective effect of BNP.

In this study, we first showed that BNP treatment increases vascularisation and the number of endothelial cells in both the ZI+BZ and RZ of infarcted hearts. Second, BNP stimulates the proliferation of endogenous pre-existing endothelial cells, likely via NPR-A binding and p38 MAP kinase activation. Third, BNP stimulates and/or accelerates the re-expression of the WT1 transcription factor in cardiac cells after MI. Fourth, in the infarcted area of untreated injured hearts, WT1^+^ EPDCs proliferate and modestly contribute to heart neovascularisation by differentiating into endothelial cells. Lastly, BNP stimulates WT1^+^ EPDC proliferation, with more endothelial cells originating from WT1^+^ EPDCs in BNP-treated infarcted hearts.

This is the first work to demonstrate that intraperitoneal injections of BNP increase neovascularisation in infarcted hearts. Thus, part of the cardioprotective effect of BNP in infarcted hearts is probably due to accelerated and/or increased neovascularisation in both the ZI+BZ and RZ. This result corroborates works reporting that natriuretic peptides stimulate angiogenesis and vasculogenesis during development and in adult ischemic organs ^33, 34^. ANP via NPR-A stimulates endothelial precursor cell proliferation, migration, and differentiation in the skin during cutaneous wound healing ^35^. The restoration of blood flow is impaired after hindlimb ischemia in NPR-A KO mice ^36^, while the injection of a high concentration of CNP increases vascular density in infarcted swine hearts ^37^. Recently, CNP was identified as a key regulator in angiogenesis and vascular remodelling after ischemia in patients suffering from peripheral artery disease ^38^. Other findings reported that BNP injections in mice lead to increased vascular regeneration in ischemic limbs ^39^ and that intramyocardial BNP gene delivery via adenovirus increased capillary density in normal rat hearts but not in infarcted hearts ^24^. Our article makes a novel contribution by showing the “time- and area-dependent” effect of BNP and the identification of both different mechanisms by which BNP increases the number of endothelial cells in infarcted hearts.

In our mouse model, the timing and action mechanisms of BNP stimulation differed in the ZI+BZ and RZ of infarcted hearts. In the RZ of infarcted hearts, the number of endothelial cells and vascularisation increased 3 days after MI. In the ZI+BZ, we detected more endothelial cells 3 days after MI but vascularisation increased only 10 days after MI. BNP stimulates endothelial cell proliferation first in the RZ (1-3 days after MI induction) and then in the ZI+BZ (10 days after MI). Although BNP seems to induce the re-expression of the WT1 transcription factor in cardiac cells in both areas of infarcted hearts (3 to 10 days after MI), its effect on WT1^+^ EPDCs is limited to ZI+BZ of infarcted hearts 10 days after MI.

The delay in stimulating endothelial cell proliferation in the different areas of infarcted hearts can be explained by the bioavailability of BNP (less cell deaths, more vessels and capillaries in the RZ), but its effect on WT1^+^ EPDCs must be favoured by the microenvironment and probably hypoxia. Indeed, in the ZI+BZ, cells undergo a synergistic effect of hypoxia and BNP.

Concerning the mechanisms, we showed that BNP acts directly on mature endothelial cells by stimulating their proliferation. The fact that BNP treatment did not increase the number of endothelial cells in neonatal NMCs isolated from neonatal NPR-A deficient hearts in our study demonstrates that BNP acts via NPR-A on neonatal cardiac cells. In the mouse model of hindlimb ischaemia, proliferating satellite cells secreted BNP, which stimulates angiogenesis in the neighbouring endothelium cells ^36^. This mechanism is also impaired in NPR-A KO mice, suggesting that regardless of the organ, BNP stimulates mature endothelial cell proliferation via the NPR-A receptor.

In our work, BNP treatment modulated also the fate of immature cells such as WT1^+^ EPDCs. This is not the first work to report the effect of BNP on immature or precursor cells. BNP level is highly correlated with the number of circulating endothelial precursor cells in patients suffering from heart failure ^39^. *In vitro*, the proliferation, adhesion, and migration capacities of endothelial precursor cells increased in a dose-dependent manner after BNP treatment ^39^. Furthermore, BNP stimulated the proliferation of satellite cells in a hindlimb ischemia model, which produced endogenous BNP stimulating the regeneration of endothelial cells ^36^.

Among cells expressing the WT1 transcription factor, it is important to discriminate between true WT1^+^ EPDCs and cardiac cells re-expressing the WT1 transcription factor in ischemic hearts ^12^. While the function of WT1 re-expression in infarcted hearts is poorly understood, it seems necessary for endothelial cell proliferation ^12^, with our results showing that BNP treatment increases and/or accelerates this process. However, in Cdh5:ROSA infarcted hearts, we detected GFP^+^ WT1^+^ BrdU^+^ cells (i.e., endothelial proliferating cells expressing WT1) as well as GFP^+^ WT1^-^ BrdU^+^ cells (i.e., endothelial proliferating cells without WT1 expression), which shows that WT1 expression is transient or unnecessary for endothelial cell proliferation.

Regarding WT1^+^ precursor cells, we induced MI in WT1:ROSA hearts and analysed the fate of GFP^+^ cells. We observed the proliferation of WT1^+^ EPDCs, mainly localised in the subepicardial layer after MI as previously reported ^12-14^. Although the differentiation of WT1^+^ cells into endothelial cells in infarcted hearts is still debated ^8, 14, 15^, we found that 5% of endothelial cells in the ZI+BZ originated from WT1^+^ EPDCs in saline-treated infarcted hearts. Interestingly, 50% of WT1^+^ cells differentiated into endothelial cells in these infarcted hearts. This shows that WT1^+^ EPDCs have the natural capacity to differentiate into endothelial cells in adult hypoxic hearts, as is the case during embryogenesis ^8^. Due their low number, it was nevertheless difficult to highlight this process.

BNP increased the number of WT1^+^ EPDCs in the subepicardial layer by stimulating their proliferation. Consequently, more cells migrated into the myocardium and differentiated into endothelial cells. We cannot exclude that BNP could also stimulate the migration of EPDCs into the myocardium, where the conditions could be adequate to differentiate into endothelial cells. Regardless of the mechanism, 10% of endothelial cells in the ZI+BZ originate from WT1^+^ EPDCs in BNP-treated hearts. However, the differentiation capacity of the WT1^+^ cells into endothelial cells was identical in saline- and BNP-treated hearts (about 50%). BNP therefore stimulates WT1^+^ EPDC proliferation but not their differentiation into endothelial cells.

However, our study does not fully elucidate the signalling pathways by which BNP induces endothelial cell and WT1^+^ EPDC proliferation. Additional work is needed in this respect, especially regarding the link between BNP treatment and VEGF expression.

In our experiment, we detected increased VEGF-A mRNA levels in the ZI+BZ and RZ of BNP-treated hearts but only 10 days after MI. We did not detect increased VEGF mRNA levels in the RZ of infarcted hearts 1 or 3 days after MI, where the proliferation of endothelial cells increased. This is consistent with the results of Kuhn et al using a hindlimb ischemia model, suggesting that BNP can promote endothelial cell proliferation independently of VEGF secretion ^36^. Thus, BNP can replace the VEGF in a non-hypoxic microenvironment to stimulate angiogenesis.

10 days after MI, in the ZI+BZ, increased mRNA levels of VEGF can result from hypoxia (i.e., from the transcriptional activity of hypoxia-inducible factor 1 alpha) and/or from BNP injections. In the RZ, higher VEGF mRNA levels were probably only linked to BNP injections 10 days after MI. As some studies reported that natriuretic peptides repress VEGF synthesis ^40^, it is unclear which mechanism(s) increased VEGF mRNA levels in both areas of the infarcted hearts 10 days after MI.

In the ZI+BZ, however, increased VEGF mRNA levels seem to induce the stimulation of WT1^+^ cell proliferation. Indeed, by stimulating WT1^+^ EPDCs with BNP, we obtained the same results as Zangi et al., who injected synthetic modified RNA encoding VEGF-A directly into the myocardium of infarcted mouse hearts, which stimulated WT1^+^ cell proliferation and shifted their differentiation into endothelial cells (i.e., 50% of WT1^+^ cells differentiated into endothelial cells) ^41^. Thus, it seems that hypoxia and BNP increase VEGF, which stimulates WT1^+^ EPDC proliferation. The interactions between BNP and VEGF should therefore be studied in more detail to confirm whether BNP acts on endothelial cells via a VEGF-independent mechanism and on WT1^+^ EPDCs via a VEGF-dependent mechanism.

The findings presented in our study may lead to new therapeutic strategies to improve the neovascularisation of hearts after ischemia. Indeed, we identified BNP as a “new cardiac angiogenic” factor, which stimulates the proliferation of both resident cardiac mature endothelial cells and WT1^+^ EPDCs. We also identified the proliferation and differentiation of WT1^+^ EPDCs as a mechanism, which can be targeted in infarcted adult hearts to increase heart revascularisation. To summarise, BNP binds to NPR-A to stimulate endothelial cell proliferation via p38 MAP kinase activation (**Fig. 10**). BNP treatment also stimulates WT1^+^ EPDC proliferation. It remains to be determined whether BNP acts directly on these cells or via increased VEGF level (**Fig. 10**). Interestingly, the benefit of LCZ696 treatment which reduces significantly the mortality of patients with chronic heart failure seems also to be associated with increased heart vascularization. However, whether this is associated with increased level of BNP remains to be demonstrated.

**Figure 10.**
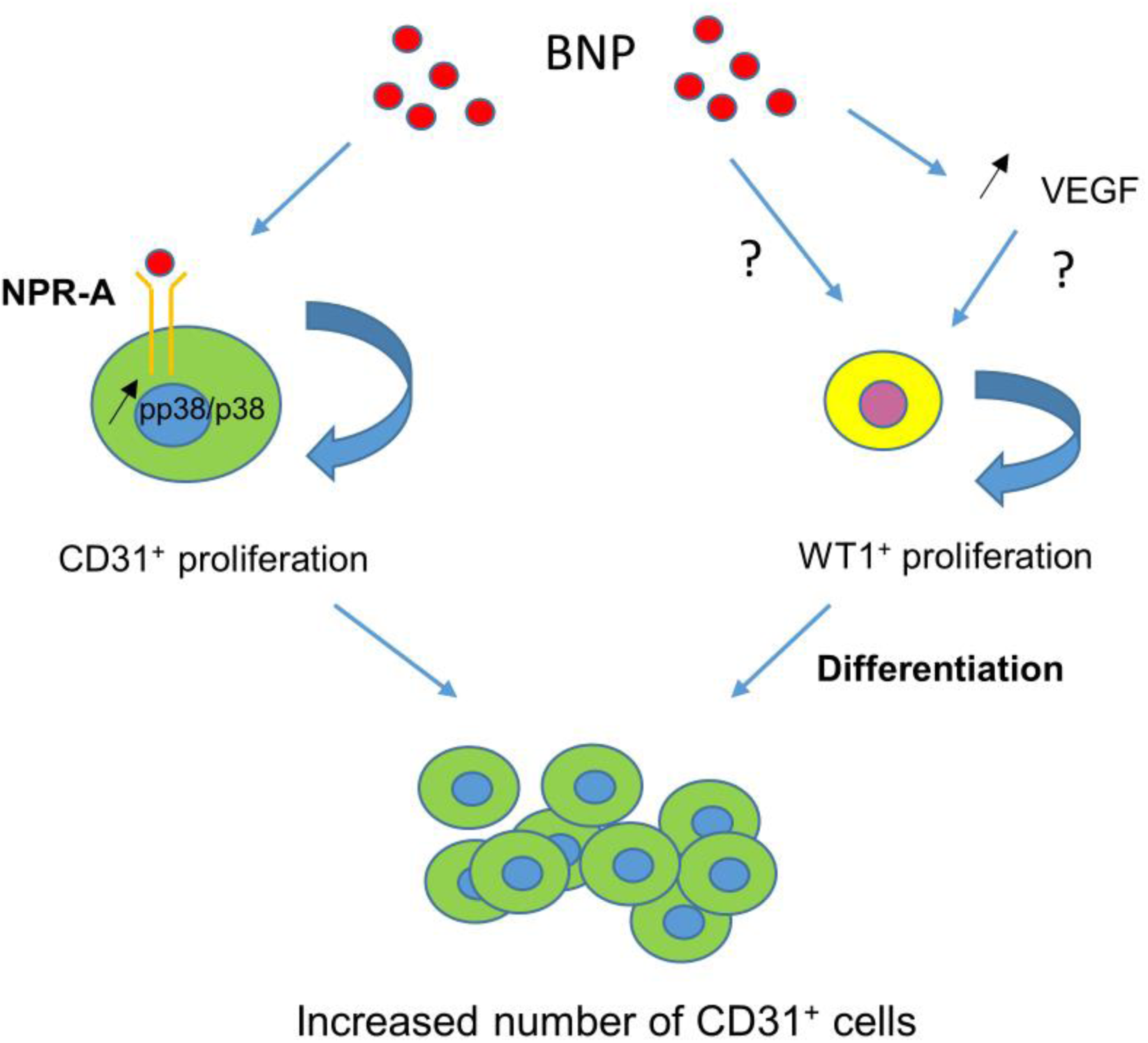
BNP-mediated mechanisms leading to increased number of endothelial cells in infarcted hearts. BNP binds directly on NPR-A receptor expressed on endothelial cells and activates p38 MAP kinase to induce their proliferation (left). BNP treatment activates also WT1^+^ EPDC proliferation in the ZI+BZ, either directly or via VEGF increase (right).

## MATERIAL AND METHODS

### Mice

All animals were maintained in accordance with the recommendations of the U.S. National Institutes of Health Guide for the Care and Use of Laboratory Animals (National Institutes of Health publication 86-23, 1985). The experiments were approved by the Swiss animal welfare authorities (authorisations VD3111 and VD3292).

C57BL/6 mice (Wild Type mice, WT) were purchased from Janvier (Le Genest-Saint-Isle, France). The NPR-A (−/−) mice were kindly provided by Dr Feng Li and Prof Nobuyo Maeda (Chapel Hill, North Carolina, US). The NPR-B deficient mouse strain (C57BL/6J-Npr2^slw^) was generated as heterozygous mice in the Laboratory of Animal Resource Bank at National Institute of Biomedical Innovation (Osaka, Japan) ^16, 26^. FUCCI (Fluorescence Ubiquitin Cell Cycle Indicator) mice were provided by Prof Marlene Knobloch ^29^. Inducible (via Tamoxifen) Cre expression under the VE Cadherin gene promoter or Cdh5(PAC)-CreER^T2^ mice were provided by Prof Tatiana Petrova ^42^. Inducible (via Tamoxifen) Cre expression under the Wilms’ tumor 1 homolog gene promoter or Wt1^tm2(cre/ERT2)wtp^/J mice were provided by Prof Thierry Pedrazzini ^43^. Gt(ROSA)26Sor^tm4(ACTB-tdTomato,-EGFP)Luo^ mice were purchased from the Jackson Laboratory (JAK 7576). Inducible Cdh5:ROSA26 and Wt1:ROSA26 mice were generated by cross-breeding Cdh5(PAC)-CreERT2 or Wt1^tm2(cre/ERT2)Wtp^/J mice with Gt(ROSA)26Sor^tm4(ACTB-tdTomato,-EGFP)Luo^ mice. All colonies were established in our animal facility.

### Experimental procedures

Only male mice were used. Myocardial infarction was induced in 8-week-old male C57 BL/6 mice by ligation of the left anterior descending coronary artery (LAD). Briefly, mice were anaesthetised (ketamine (65 mg/kg)/xylazine (15 mg/kg), acépromazine (2 mg/kg)), intubated and ventilated. The chest cavity was entered through the third intercostal space at the left upper sternal border, and myocardial infarction was induced by ligature of the LAD with a 7-0 nylon suture at about 1–2 mm from the atria. Sham-operated mice underwent the same operation but without tightening the knot.

For Cdh5:ROSA and Wt1:ROSA mice, surgeries were performed in 8 week-old adult male mice injected 2 weeks before with Tamoxifen as previously described. Tamoxifen (Sigma, T5648) was dissolved in ethanol at 100 mg/ml and emulsified in peanut oil to a final concentration of 10 mg/ml. 1 mg Tamoxifen/25g body weight was injected intraperitoneally (i.p.) to adult mice every 3 days for 3 times ^16^.

Directly after the surgery, NaCl or BNP (1 µg / 20 g mouse in 20 µl, Bachem synthetic mouse BNP (1-45) peptide (catalog number H-7558)) was injected into the left ventricle cavity. Temgesic (Buprenorphine, 0.1 mg/kg) was injected subcutaneously as soon as the mice waked up and every 8-12 hours during 2 days.

BNP (2 µg / 25 g mouse) was injected i.p. every 2 days ^16^. After surgery (for the mice sacrificed after 1 day) or 24h after the surgery (for all other mice), BrdU (1 mg/ml, Sigma B5002) was added to drinking water and changed every 2 days during 10 days.

For the experiments related to LCZ696 (Entresto, Novartis) treatment, mice after myocardial infarction were randomly assigned into 3 different groups: H2O, LCZ696 [6 mg/kg/day], or LCZ696 [60 mg/kg/day]. These two drug concentrations were chosen as LCZ696 [6 mg/kg/day] is the dose mostly used in patients (200-600 mg/day) and LCZ696 [60 mg/kg/day] induces a dose-dependent increase in plasmatic natriuretic peptides in animals ^44^. LCZ696 drugs were grounded, formulated in water and sonicated for 1 hour before administration. Drugs were administrated 24 hours after MI and once daily for 10 days by oral gavage. Blood pressure was measured daily, from one week before surgery until sacrifice using a tail-cuff based CODA high throughput system (Kent Scientific Corporation).

Mice were sacrificed 1, 3, 10, or 28 days after infarct induction and hearts were removed (Fig. 1A). If immunofluorescence has to be carried out, apex was embedded into OCT and slowly frozen. Remaining heart was separated into 3 zones, the infarct zone (ZI), the border zone (BZ) and the remote zone (RZ). According the required experiment, ZI and BZ may be pooled. Tissues were either digested for flow cytometry analysis or quickly frozen for mRNA or protein analysis.

### Cell culture

Non-myocyte cells (NMCs) were isolated from the hearts of neonatal C57BL/6, NPR-A KO or NPR-B KO pups (1-2 days) as previously described ^16, 26^ and were cultured in medium composed of MEM Alpha (Gibco 32571-028), 10% FBS, 100 U/ml penicillin G, 100 µg/ml streptomycin with or without BNP (5 µg/ml) up to confluence (i.e., 10-11 days). Adult cardiac NMCs were isolated from adult C56BL/6 mice (6-8 weeks old) by digesting adult ventricles in buffer containing 1 mg/ml collagenase IV (Gibco 17104-019) and 1.2 mg/ml dispase II (Sigma, D4693) and were cultured in EGM™-2 Endothelial Cell Growth Medium-2 BulletKit (Lonza, CC-3162) supplemented with 15% fetal calf serum (FCS) (invitrogen Corp) with or without BNP (5 µg/ml) up to confluence (i.e., 5-10 days). Neonatal and adult NMCs were maintained at 37°C in 5% CO_2_ and 3% O_2_.

### Flow cytometry analysis

Cultured neonatal or adult NMCs were removed from dishes using Cell dissociation buffer enzyme-free PBS-based (Gibco 13151-014) and washed in PBS with 3% fetal calf serum. Adult NMCs were isolated from adult infarcted hearts as described above. Samples were treated 5 minutes at room temperature using CD16/CD32 antibody (BD Biosciences, 553142, 1μl/10^6^ cells) and stained with different antibodies listed in Supplemental Table S1 (Supplemental Informations). In case of WT1:ROSA26 cells, cytoplasmic staining was done for GFP detection after fixation (1.5% PFA) and permeabilisation (0.2% saponin). All stainings were performed 20 min on ice. Cells were analyzed with Gallios cytometer and data using FlowJo 10 software.

The numbers of CD31^+^ cells in cell cultures or in NMCs isolated from hearts were obtained by relating the percentage of the CD31^+^ cells obtained by flow cytometry analysis and the total number of NMCs in culture or obtained after heart digestion in the different area of infarcted hearts. The number of GFP^+^ or GFP^-^ cells among NMCs isolated from infarcted Cdh5:ROSA mice was determined by the same method, using flow cytometry analysis.

### Endothelial Cell sorting

NMCs were isolated from Cdh5:ROSA hearts, injected with Tamoxifen 2 weeks before. GFP^+^ cells were sorted with the MoFlo Astrios Flow Cytometer System (Beckman Coulter). Then cells were split in half and treated or not with BNP (5 μg/ml) for 1h30 at room temperature. Cells were lysed and proteins extracted.

### Echocardiography and measurements

Transthoracic echocardiographies were performed on adult unmanipulated or infarcted mice using a 30 M-Hz probe and the Vevo 770 Ultrasound machine (VisualSonics, Toronto, Ontario, Canada) as described ^16^. All measurements were done from leading edge to leading edge according to the American Society of Echocardiography guidelines. Ejection fraction (EF) and fractional shortening, were evaluated on lightly anaesthetised mice (1 % isoflurane). Furthermore, according to the fact that changes in left ventricle volume can be considered as an index of remodeling ^45^, we calculated the percentage of increase of the left ventricle volume 10 days after surgery, which is the ratio between (LV Vol;d 1 or 4 weeks − LV Vol;d before surgery) and LV Vol;d before surgery × 100.

### Statistical analysis

All results were presented as mean ± SEM. Statistical analyses were performed only if the number of experiments or mice is ≥ 6 per group. The paired or unpaired Student-T test or Wilcoxon-Mann-Whitney test were used. The alpha level of all tests was 0.05.

**Please refer to the supplementary file for more informations about the methods, primers and antibodies used.**

## Supporting information

Supplemental informations

## ACKNOWLEDGMENTS

The authors thank Dre Corinne Bertonneche and Ms Anne-Catherine Clerc for her technical expertise.

This research was funded by the Swiss National Foundation (PMPDB 310030_162985), the Novartis Foundation for medical-biological research and the Emma Muschamp Foundation (Lausanne).

## AUTHOR CONTRIBUTIONS

**Na Li:** Formal analysis, Investigation, Methodology, Project administration, validation, visualisation

**Stéphanie Clerc-Rignault:** Data curation, formal analysis, investigation, methodology, resources

**Christelle Bielmann:** Formal analysis, investigation, methodology

**Anne-Charlotte Bon-Mathier:** Investigation, resources, methodology, formal analysis

**Tamara Déglise:** Formal analysis, investigation, methodology

**Alexia Carboni and Mégane Ducrest:** Master students working on the LCZ696 treatment.

**Nathalie Rosenblatt-Velin:** Conceptualisation, data curation, funding acquisition, supervision, validation, writing-original draft.

## CONVENTIONS AND ABBREVIATIONS

ANP: atrial natriuretic peptide
BNP: brain natriuretic peptide
BrdU: 5-Bromo-2′-deoxyuridine
CNP: C-type natriuretic peptide
EndMT: endothelial-to-mesenchymal transition
eNOS: endothelial nitric oxide synthase
EPDC: epicardium-derived cell
FUCCI: fluorescent ubiquitination-based cell cycle indicator
GFP: green fluorescent protein
HUVEC: human umbilical vein endothelial cells
KO: knockout mouse model
MI: myocardial infarction
NMC: non-myocyte cells
NPR-A and NPR-B: natriuretic peptide receptor A and B
RZ: remote zone
Sca-1: stem cell antigen 1
SMA: smooth muscle actin
VEGF: vascular endothelial growth factor
WT1: Wilms’ tumour 1 transcription factor
ZI+BZ: infarct zone and border zone

