## Supplemental informations for "Increasing heart vascularisation after myocardial infarction using brain natriuretic peptide stimulation of endothelial and WT1^+^ epicardium-derived cells"

##### Detailed methods

**Supplemental Table S1.** Antibodies used in flow cytometry analysis, immunohistology and Western blot analysis.

**Supplemental Table S2:** Primer Sequences used in quantitative RT-PCR.

### Detailed Methods

##### *Immunofluorescence*

Neonatal and adult hearts were embedded in OCT. Immunostainings were performed on 5µm heart sections or on cells cultured for up to 11 days on coverslips. Tissue sections or cells were fixed 10 min in 2% PFA. The first antibodies were all incubated overnight at 4 °C. Secondary antibodies were incubated 1 h at room temperature (Supplemental Table S1). For BrdU detection, heart slides were fixed 10 minutes in 2% PFA, DNA was denaturated 1h at room temperature in HCl 2N before neutralisation in Na Borate 0.1M pH=8.5, 2x5 min. Rat anti-BrdU (1/100, Abcam) was incubated 1h at room temperature. Donkey anti-rat was used as secondary antibody. Nuclei were stained with DAPI (0.3 µM). All slides were mounted with Dabco mounting medium (Sigma D2, 780-2) and examined with a Nikon eclipse 90i microscope or Nikon SMZ 25 Stereomicroscope (for the hearts in full, Figure 8).

To note, no GFP staining was required to detect the GFP positive cells in the Tamoxifen injected Cdh5:ROSA mice. However, we used an antibody anti-GFP to detect the $GFP^+$ cells in the WT1:ROSA mice injected with Tamoxifen.

To study heart vascularization, the number of pixels was obtained by processing immunostaining pictures with Adobe Photoshop software.

The percentages of proliferating endothelial ($CD31^+$) or $WT1^+$ cells were obtained by dividing the number of $CD31^+$ $BrdU^+$ cells or $WT1^+BrdU^+$ cells obtained by counting per the total number of $CD31^+$ cells or $WT1^+$ cells, respectively.

##### *Quantitative RT-PCR*

Total RNA was isolated from heart tissue or cell culture using Trizol (Ambion 15596026). Reverse transcriptase was carried out using PrimeScript™ RT Reagent kit with gDNA eraser (perfect Real Time) (Takara, RR047A).

Quantitative real time polymerase chain reaction was performed in duplicates using the TB Green™ Premix Ex Taq™ kit (Takara RR420L) on a ViiA™ 7 Instrument (Applied Biosystems). Results were obtained after 40 cycles of a thermal step protocol consisting of 95 °C (1 s), 60 °C (20 s). The primer sequences were reported in Supplemental Table S2. Gene expressions were normalized using the housekeeping gene 18S (ΔCT values). Means of ΔΔCT values (versus untreated cells) were calculated and results were represented as $2^{-\Delta\Delta CT}$. Statistics were performed on ΔΔCT individual values [2].

***Western blot***

Total proteins were extracted from tissues or cells as described [1, 2] and transferred to nitrocellulose membranes before incubation with primary antibodies overnight at 4°C (Supplemental Table S1). Secondary antibodies were added 2 hours at room temperature. The immunoblot signals were detected and quantified using the Odyssey infrared imaging system (LI-COR Biosciences, Bad Homburg, Germany). All results were related to their expression of tubulin.

***Determination of cGMP concentration in plasma***

cGMP level was detected using the cGMP Enzyme Immuno Assay kit Direct (Sigma). BNP was injected in unmanipulated or infarcted mice. Blood was collected 1-2h after BNP injection for unmanipulated mice and 1or 3 days after surgery for infarcted hearts. EDTA-plasma were then processed as recommended in the kit.

**Supplemental Table S1.** Antibodies used in flow cytometry analysis, immunohistology and Western blot analysis.

| Antibodies | Species | Dilution | Reference | usage |
|---|---|---|---|---|
| **anti-goat Alexa 488** | chicken | 1/500 | Molecular Probes A21467 | Flow cytometry |
| **anti-rabbit Alexa 488** | donkey | 1/1000 | Molecular Probes A21206 | Flow cytometry |
| **CD31 Biotin** | rat | 1/50 | BD Biosciences 553371 | Flow cytometry |
| **GFP** | rabbit | 1/500 | Abcam ab290 | Flow cytometry |
| **Isotype control for CD31** | rat | 1/10 | BD Biosciences 553928 | Flow cytometry |
| **NPRA** | rabbit | 1/50 | Abcam ab70848 | Flow cytometry |
| **NPRB** | goat | 1/50 | Santa Cruz Sc-34421 | Flow cytometry |
| **Streptavidine APC** | rat | 1/1000 | BioLegend 405207 | Flow cytometry |
| **anti-goat Alexa 594** | donkey | 1/1000 | Molecular Probes A11058 | immunohistology |
| **anti-goat biotin** | horse | 1/200 | Vector BA-5000 | immunohistology |
| **anti-rabbit Alexa 488** | goat | 1/1000 | Molecular Probe A11034 | immunohistology |
| **anti-rabbit Alexa 488** | donkey | 1/1000 | Molecular Probes A21206 | immunohistology |
| **anti-rabbit Alexa 594** | donkey | 1/1000 | Molecular Probes A21207 | immunohistology |
| **anti-rat Alexa 488** | donkey | 1/1000 | Molecular Probes A21208 | immunohistology |
| **anti-rat Alexa 647** | donkey | 1/1000 | Jackson Immuno 712-605-150 | immunohistology |
| **BrdU** | rat | 1/100 | Abcam ab6326 | immunohistology |
| **CD31** | rabbit | 1/100 | Abcam ab28364 | immunohistology |
| **CD31 biotin** | rat | 1/100 | BD Biosciences 553371 | immunohistology |
| **CD45** | rabbit | 1/50 | Abcam ab10558 | immunohistology |
| **ckit** | rat | 1/100 | R&D Sytems MAB1356 | immunohistology |
| **GFP** | goat | 1/2000 | Abcam ab5450 | immunohistology |
| **NPRA** | rat | 1/50 | R&D Sytems MAB3974 | immunohistology |
| **NPRA** | rabbit | 1/50 | Abcam ab70848 | immunohistology |
| **NPRB** | goat | 1/50 | Santa Cruz Sc-34421 | immunohistology |
| **NPRB** | rabbit | 1/20 | Abcam ab139188 | immunohistology |
| **Sca-1** | rat | 1/1000 | Abcam ab51317 | immunohistology |
| **Streptavidine Alexa 594** | | 1/1000 | Molecular Probe S11227 | immunohistology |
| **Streptavidine Alexa 647** | | 1/1000 | Molecular Probe S32357 | immunohistology |
| **WT-1** | rabbit | 1/100 | Abcam ab89901 | immunohistology |
| **anti-mouse IRDye 800** | goat | 1/10000 | Rockland 610-132-121 | Western Blot |
| **anti-rabbit Alexa 680** | goat | 1/5000 | Molecular Probe A21109 | Western Blot |
| **CD31** | rabbit | 1/500 | Abcam ab28364 | Western Blot |
| **Phospholamban** | rabbit | 1/1000 | Millipore | Western blot |
| **Phospho phospholamban** | mouse | 1/500 | Millipore | Western blot |
| **Phospho p38** | rabbit | 1/500 | Cell Signaling | Western Blot |
| **p38** | rabbit | 1/1000 | Cell Signaling | Western blot |
| **tubulin** | mouse | 1/10000 | Sigma T5168 | Western Blot |

**<u>Supplemental Table S2:</u>** Primer Sequences used in quantitative RT-PCR.

| Gene | Sense | Anti-sense | Product size (bp) |
|---|---|---|---|
| **18S** | ACTTTTGGGGCCTTCGTGTC | GCCCAGAGACTCATTTCTTCTTG | 96 |
| **alpha SMA** | CAGGCATGGATGGCATCAATCAC | ACTCTAGCTGTGAAGTCAGTGTCG | 154 |
| **CD31** | GCCTCACCAAGAGAACGGAAGGC | CTGCTTTCGGTGGGGACAGGC | 158 |
| **CD34** | CTTCTGCTCCGAGTGCCATT | GCCAAGACCATCAGCAAACAC | 250 |
| **c-kit** | ATCTGCTCTGCGTCCTGTTG | CTGATTGTGCTGGATGGATG | 108 |
| **eNOS** | GGCTGTGGTAGTTAGGGCATC | AGGTTTGGGTTGGGCATCT | 165 |
| **Flk1** | ACTGCAGTGATTGCCATGTTCT | CCTTCATTGGCCCGCTTAA | 74 |
| **Sca-1** | TTTGAGACTTCTTGCCCATC | ACCCAGGATCTCCATACTTTC | 159 |
| **Ve-cad** | AGCGCAGCATCGGGTACT | TCGGAAGAATTGGCCTCTGT | 56 |
| **vWF** | GATGCCCCAGTCAGCTCTAC | TCAGCCTCGGACAACATAGA | 131 |
| **WT1** | CACGGCACAGGGTATGAGAG | GTTGGGGCCACTCCAGATAC | 128 |

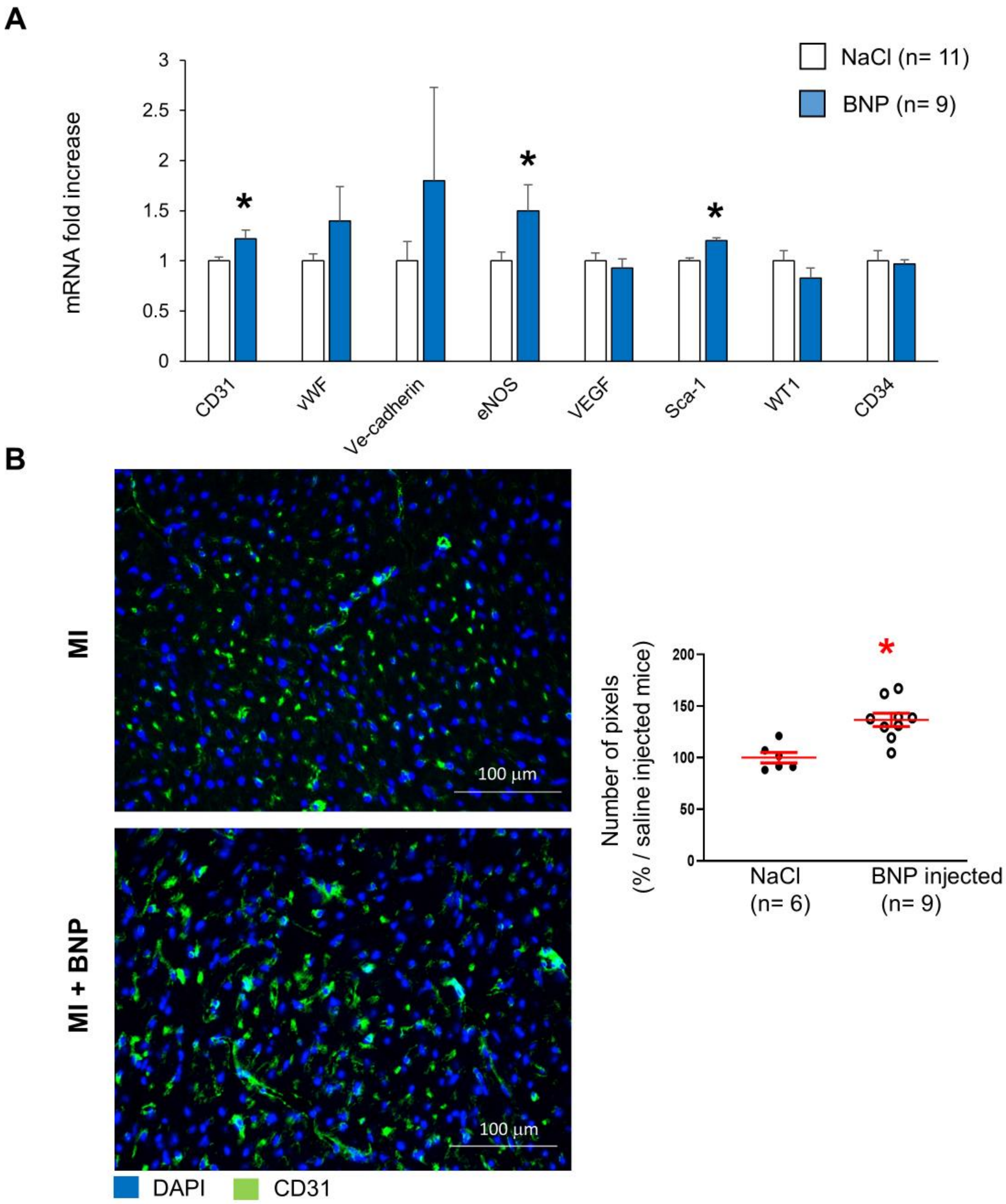


<u>Supplemental Figure 1</u>. **BNP injection led to increased vascularization in unmanipulated hearts**. **A**. Quantitative relative expression of mRNAs coding for CD31, von Willbrand factor (vWF), Ve-cadherin, eNOS, VEGF, Stem Cell antigen 1 (Sca-1), Wilms' tumor 1 (WT1) and CD34. **B.** Representative immunostainings against CD31 protein (green) of saline and BNP treated unmanipulated hearts. Nuclei stained in blue with DAPI. CD31 staining intensity measured on at least 10 different pictures per heart and per area. The numbers of pixel obtained for saline treated hearts related to the numbers obtained in BNP injected hearts. All the results are means ± SEM, * $p<0.05$.

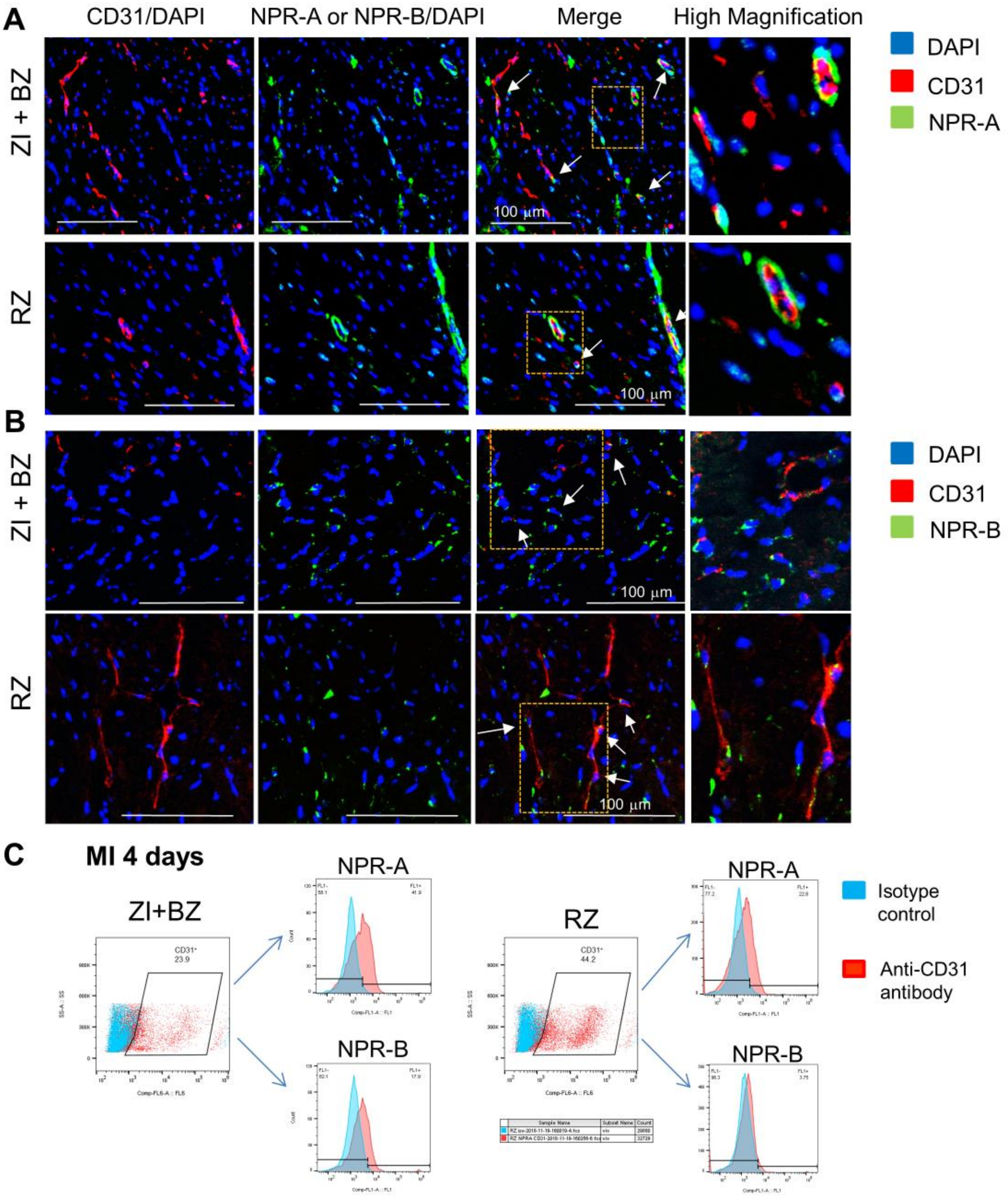


**Supplemental Figure 2. Adult cardiac endothelial cells express BNP receptors in both infarcted and border (ZI+BZ) and remote (RZ) zones.** The presence of NPR-A (**A**) or NPR-B (**B**) was assessed by immunostainings on adult hearts 24 hours after MI or by flow cytometry analysis (**C**) on adult non-myocytes cells isolated from hearts 4 days after MI. Antibody against CD31 was used in combination with anti-NPR-A or NPR-B antibodies. The percentages of NPR-A or NPR-B expressing cells were evaluated on $CD31^{+}$ cells.

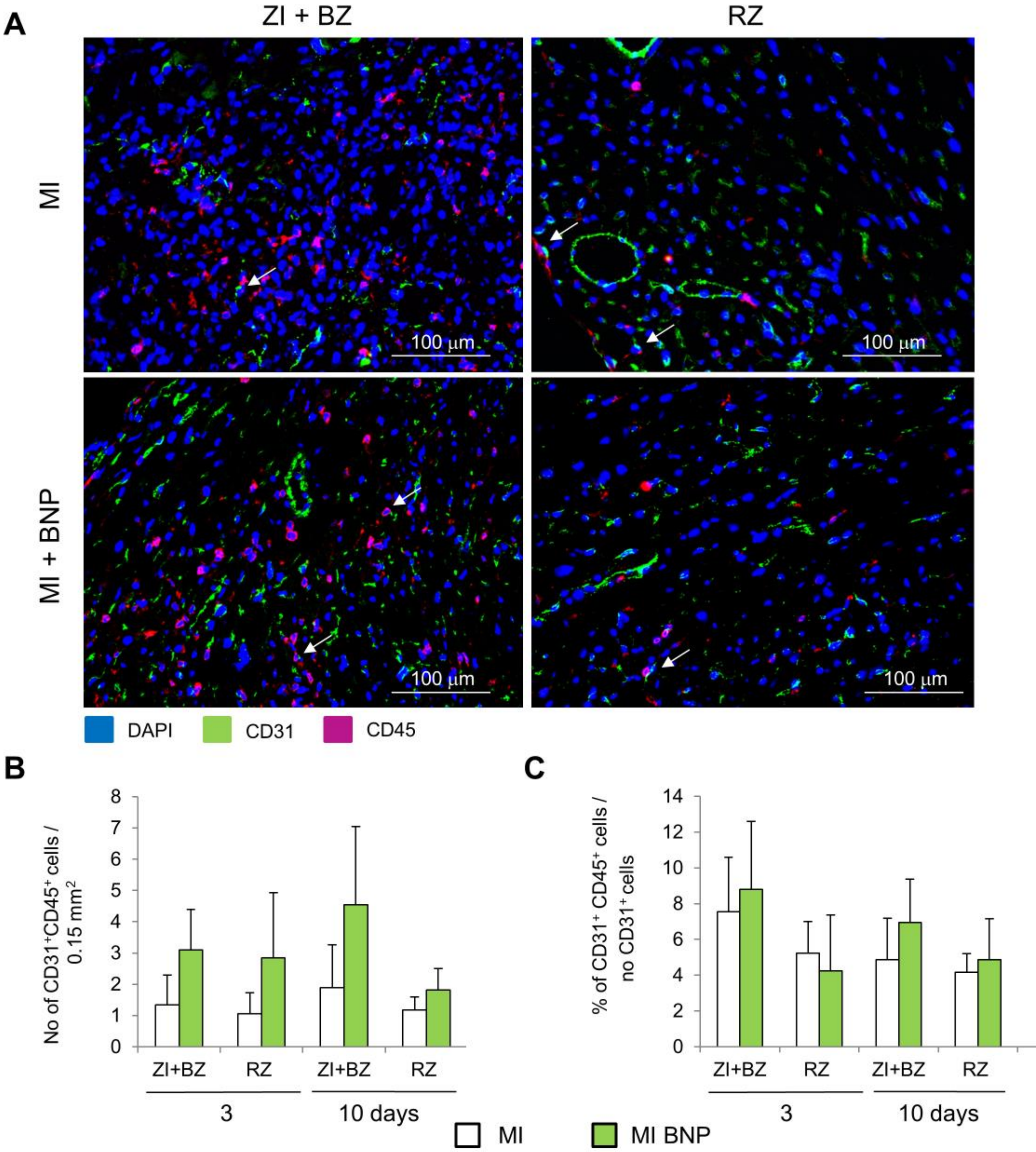


<u>Supplemental Figure 3.</u> **Few cardiac endothelial cells in infarcted and remote zone 3 and 10 days after infarction express the CD45 protein.** The presence of $CD31^+$ $CD45^+$ cells was assessed by immunostainings on adult hearts 3 and 10 days after MI. (**A**) Representative pictures of the stainings 10 days after MI. (**B**) The numbers of double positive cells were counted (**B**) on heart sections (0.15 $mm^2$) of the different area of saline (MI) and BNP treated infarcted hearts (MI + BNP) and related (**C**) to the total number of $CD31^+$ cells. Cells were counted on at least 10 different pictures per area and mouse. n= 4-6 mice per group. All the results are means ± SEM.

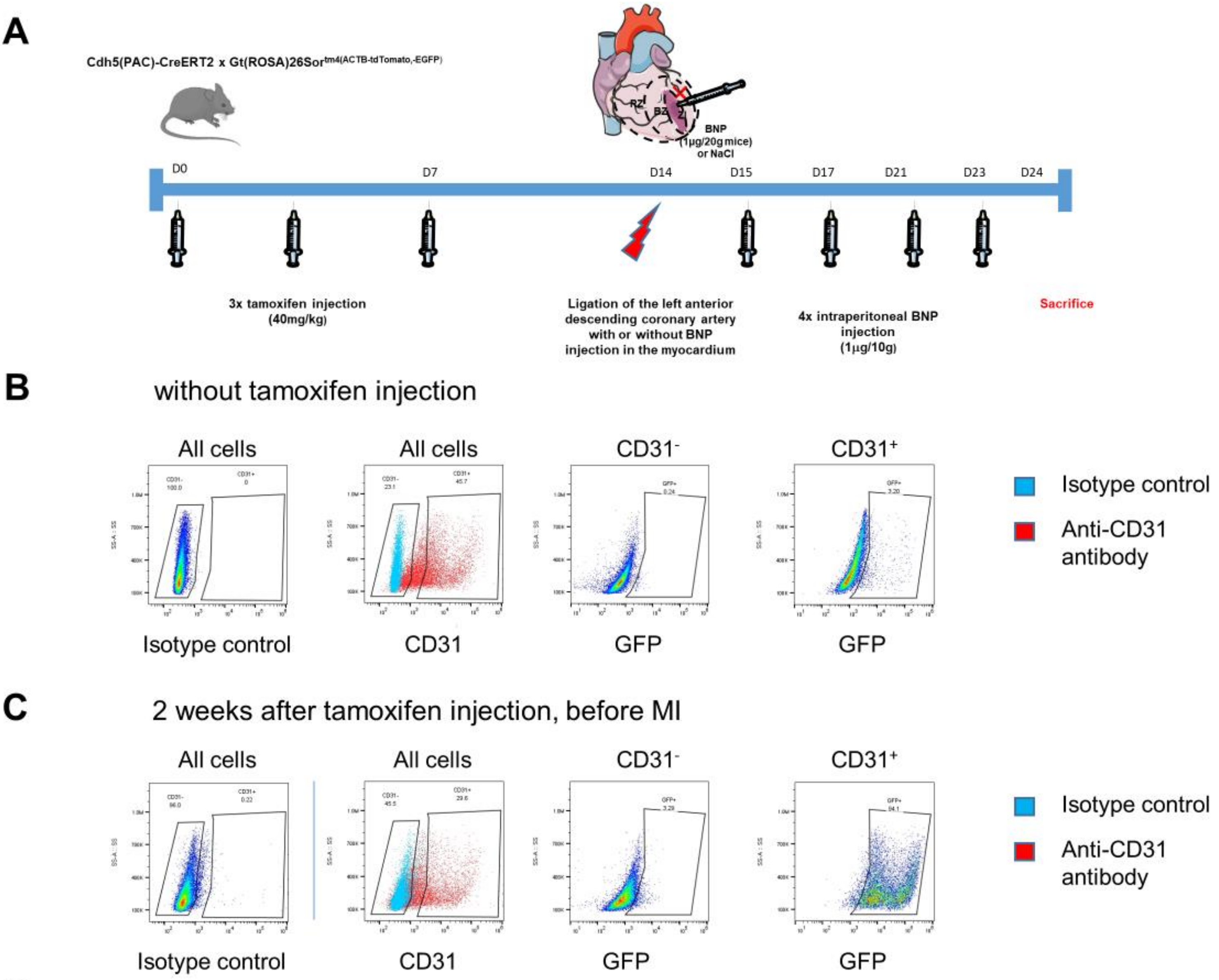


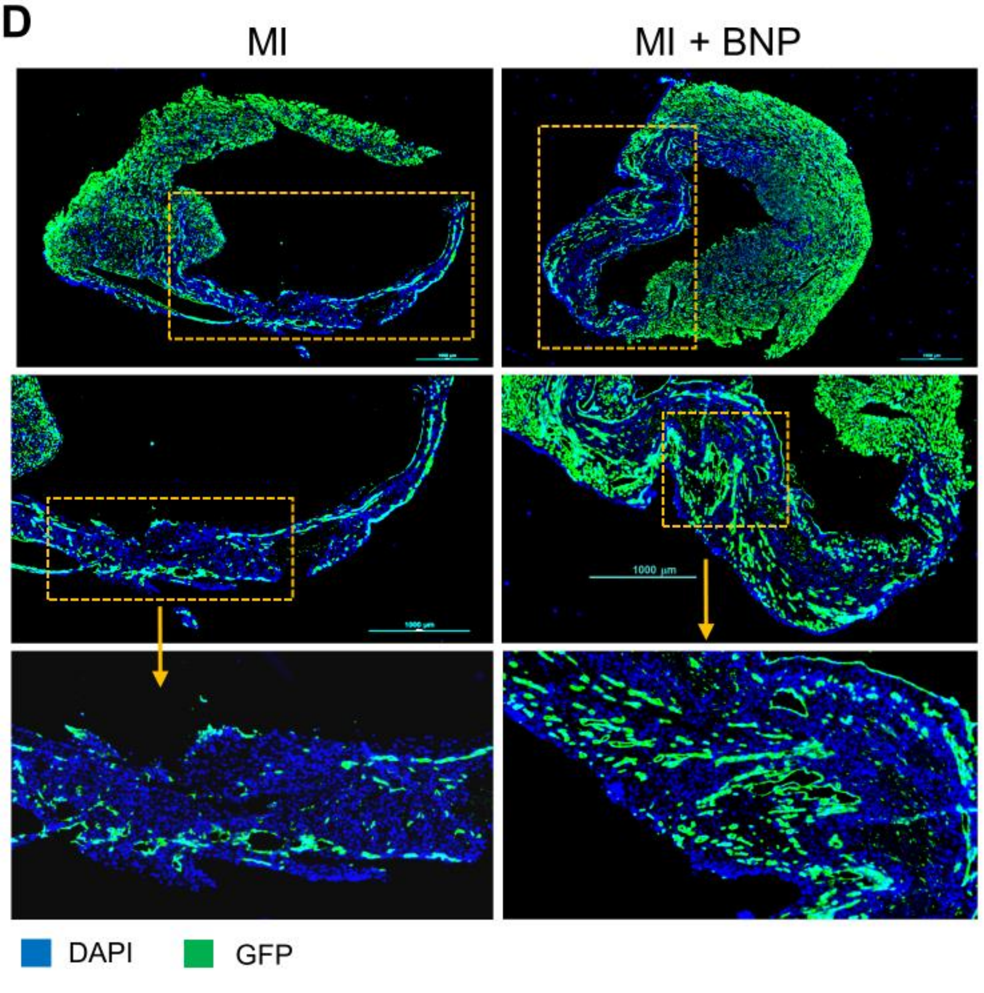


**Supplemental Figure 4. Characterisation of the Cdh5:ROSA mouse model.** **A.** Heterozygous Cdh5 Cre x ROSA mice were used to follow the Ve-Cadherin expressing cells. **B.** Without tamoxifen injection no CD31⁺ cells expressed the GFP protein. **C.** At time of surgery, 2 weeks after the first tamoxifen injection, 94% of CD31⁺ cells were GFP positive. **D.** Representative pictures of infarcted hearts 10 days after MI. Only DAPI staining. The ZI+BZ in orange rectangles are represented at high magnification.

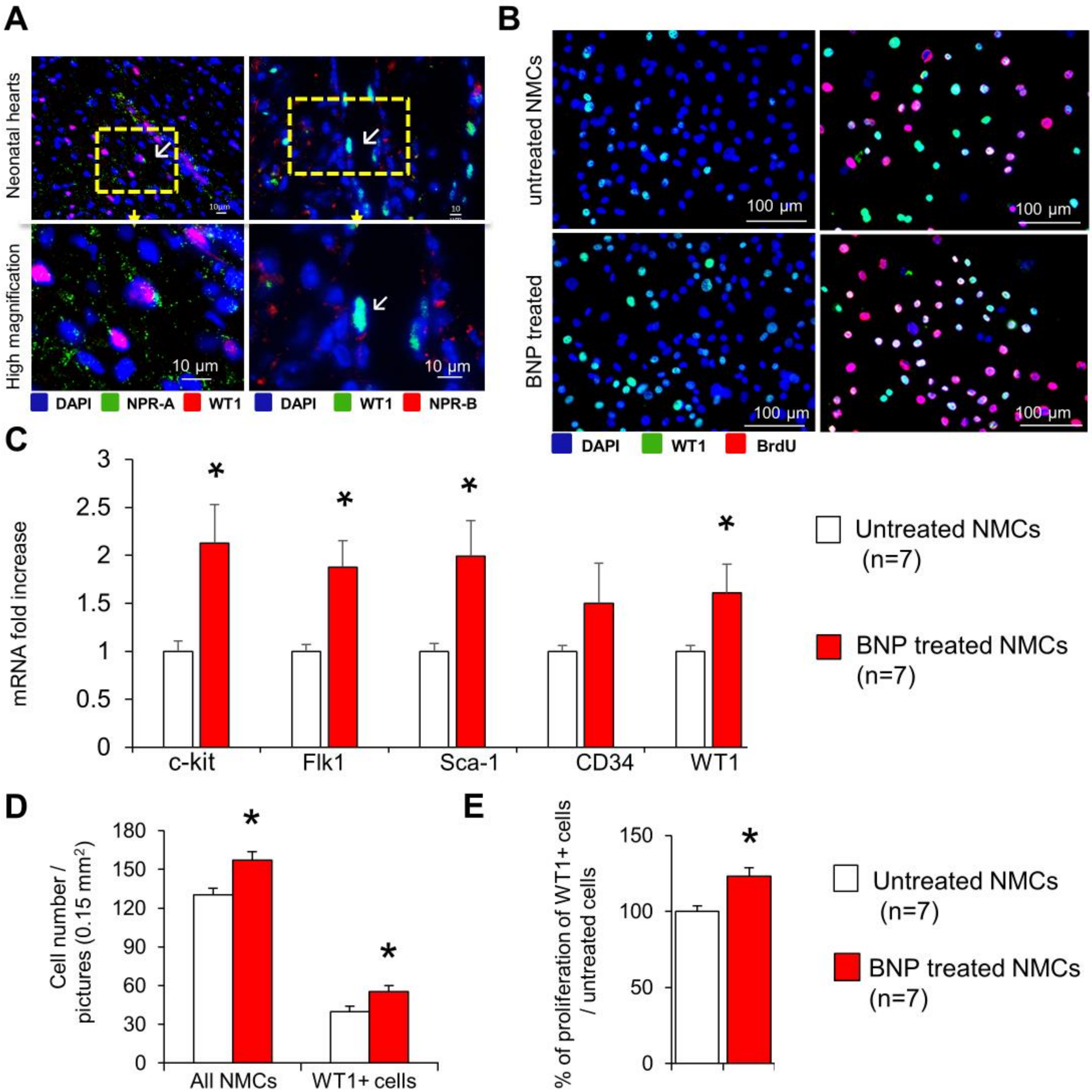


**Supplemental Figure 5. BNP treatment stimulated WT1⁺ cell proliferation *in vitro*. A.** WT1+ cells isolated from neonatal hearts express NPR-A and NPR-B. **B**. Representative immunostainings of NMC cultures treated or not with BNP (5 µg/ml) and stained with DAPI (Nuclei in blue), antibodies against WT1 (green) and BrdU (red). **C.** Quantitative relative expression of mRNAs coding for endothelial precursor specific genes (Flk-1, c-kit, stem cell antigen 1 (Sca-1), CD34 and Wilms' tumor 1 (WT1)) in NMCs isolated from neonatal hearts and treated or not with BNP (5 mg/ml) for 10 days. Results expressed as fold-increase above the levels in untreated cells. **D.** Quantification of the number of $WT1^+$ cells per pictures (0.15 $mm^2$) after 7-10 days of culture. At least 10 different pictures were evaluated per cell culture. **E**. The percentages of $WT1^+$ cell proliferation were obtained by dividing the number of $WT1^+$ $BrdU^+$ cells per the total $WT1^+$ cell number. Results expressed as fold-increase above the levels in untreated cells All the results are means ± SEM, * $p<0.05$.

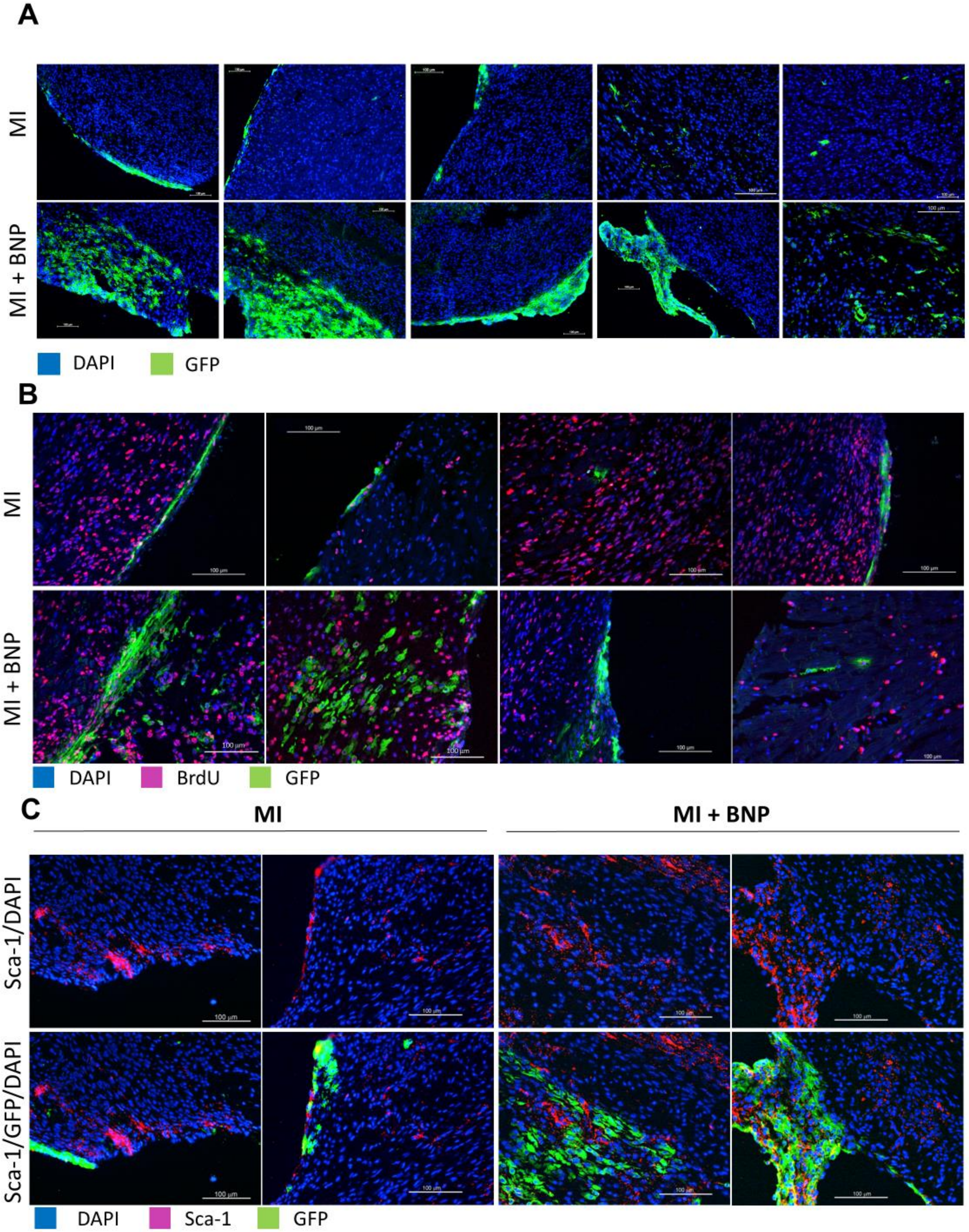


**<u>Supplemental Figure 6.</u> Representative pictures of infarcted WT1:ROSA mice treated or not with BNP, 10 days after surgery. A.** Cells expressing the GFP protein were mainly localized in the epicardium in MI hearts and migrate to the endocardium in BNP treated infarcted hearts. **B.** Some GFP$^+$ cells proliferated and/or expressed the Sca-1 protein (**C**).
